# Identification of a TNF-TNFR-like system in malaria vectors (*Anopheles stephensi*) likely to influence *Plasmodium* resistance

**DOI:** 10.1101/2022.07.28.501812

**Authors:** Subhashini Srinivasan, Chaitali Ghosh, Shrestha Das, Aditi Thakare, Siddharth Singh, Apoorva Ganesh, Harsh Mahawar, Aadhya Jaisimha, Mohanapriya Krishna, Aritra Chattopadhyay, Rishima Borah, Vikrant Singh, M Soumya, Naveen Kumar, Sampath Kumar, Sunita Swain, Suresh Subramani

## Abstract

Identification of *Plasmodium*-resistance genes in malaria vectors remains an elusive goal despite the recent availability of high-quality genomes of several mosquito vectors. *An. stephensi,* with its three distinctly-identifiable forms at the egg stage, correlating with varying vector competence, offers an ideal species to discover functional mosquito genes implicated in *Plasmodium* resistance. Recently, the genomes of several strains of *An. stephensi* of the type-form, known to display high vectorial capacity, have been reported. Here, we report a chromosomal-level assembly of an intermediate-form of *An. stephensi* strain (IndInt), shown to have reduced vectorial capacity relative to a strain of type-form (IndCh). The contig level assembly with a L50 of 4 was scaffolded into chromosomes by using the genome of IndCh as the reference. The final assembly shows a heterozygous paracentric inversion, 3L*i,* involving 8 Mbp, which is syntenic to the extensively-studied 2L*a* inversion implicated in *Plasmodium* resistance in *An. gambiae* involving 21 Mbp. Deep annotation of genes within the 3L*i* region in IndInt assembly using the state-of-the-art protein-fold prediction and other annotation tools reveals the presence of a TNF-like gene, which is the homolog of the eiger gene in *Drosophila.* Subsequent chromosome-wide searches revealed homologs of wengen (wgn) and grindelwald (grnd) genes in IndInt, which are known to be the receptors for eiger in *Drosophila*. We have identified all the genes in IndInt required for eiger-mediated signaling by analogy to TNF-TNFR system, suggesting the presence of a functionally active eiger signaling pathway present in IndInt. Comparative genomics of high-quality genome assemblies of the three type-forms with that of IndInt, reveals structurally disruptive mutations in eiger gene in all three strains of the type-form, alluding to compromised innate immunity in the type-form as the cause of high vectorial capacity in these strains. This is the first report of the presence of an intact evolutionarily-conserved TNF-TNFR signaling system in malaria vectors, with a potential role in *Plasmodium* resistance.

## INTRODUCTION

*Anopheles stephensi* is an established urban malarial vector in India and South East Asia with expanding range into African cities. This vector quickly adapts to the local environment, survives extremely high temperatures, and establishes resistance to multiple insecticides (Vector alert: *Anopheles stephensi* invasion and spread, 2019). Based on the vectorial capacity of *An. stephensi*, there are three variants classified as type-form (high vectorial capacity), intermediate-form (moderate vectorial capacity), and mysorensis (low vectorial capacity)[1] [2]. These forms are identifiable based on the number of ridges on eggs[3]. The genomes of three strains of the type-form have been reported recently[4][5][6]. A comparison of these three genomes revealed the presence of all three genotypes of the 2R*b* inversion among the various strains of the type-form.

Sixteen paracentric inversions in the chromosomes of *An. stephensi* have been reported by visualizing loop formation in chromosomes from large heterozygous inversions. While chromosome X has no reported inversions, chromosomal arms 2R, 2L, 3R, and 3L have many paracentric inversions characterized based on cytobands[7][8]. Moreover, each of these inversions are reported to impart varying resistance to insecticides[9]. However, during the maintenance of various strains of *An. stephensi* under laboratory conditions, several inversions are lost within the first few generations, such as 2R*e*, 3R*c,* and 3L*c*, whereas other inversions, such as 2R*b*, 3R*b,* and 3L*i,* persist through generations even in the absence of any environmental pressure, and in some cases, increasing in frequency with increasing generations[7]. Based on these reports and studies on other malaria vectors, it is now well established that chromosomal inversions are dynamically used by vectors to adapt to environmental changes while retaining fitness[10].

*An. gambiae,* a malaria vector from Africa, is by far the most extensively studied and interrogated using genomics. The inversions 2R*b* and 2L*a* in *An. gambiae* have been extensively characterized at the molecular and population levels. While the 2R*b* inversion in *An. gambiae,* involving 8 Mbp, is associated with resistance to DDT and adaptation to climate[10], the 2L*a* inversion, encompassing 21 Mbp, is associated with resistance to *Plasmodium* and desiccation[11][12]. The breakpoints of 2L*a* have been extensively studied by selecting BAC clones spanning breakpoints from strains with alternate arrangements of the 2L*a* configuration[13]. A systems-level study of gene expression in *An. gambiae* between different configurations of a given inversion shows that a large number of genes within the inversion loci are differentially regulated[14].

Comparative proteomics of *D. melanogaster, Ae. aegypti, and An. gambiae* revealed 91 gene trios with a total of 589 genes comprising the immune repertoire[15]. These are homologs of genes from pathways implicated in classical innate immunity or defense functions in insects. The pathways include apoptosis inhibitors (IAPs), oxidative defense enzymes [superoxide dismutases (SODs), glutathione peroxidases (GPXs), thioredoxin peroxidases (TPXs), and heme-containing peroxidases (HPXs)], and class A and B scavenger receptors (SCRs) common to all three species. Of these, more recently, homologs of 361 genes were reported in *An. stephensi*[4] (https://github.com/mahulchak/stephensi_genome/blob/master/immune.gff), which is in the same range (285, 338, and 353, respectively) of immune-repertoire genes reported for *D. melanogaster* (*Dm*), *An. gambiae* (*Ag*), and *Ae. aegypti* (*Aa*), respectively.

Deep annotation of genes within inversion regions associated with resistance to pathogens offers an opportunity to discover additional immune repertoire genes that may have diverged beyond the point of recognition by standard sequence-based annotation tools. RoseTTaFold, a state-of-the-art protein-fold prediction tool, allows deeper annotation of uncharacterized or divergent proteins, thereby aiding the identification of hidden/missed immune-related genes in *Anopheles*[16]. The idea is that the diversity in proteins far exceeds the number of folds in nature, which is estimated to be as low as one thousand[17]. This is especially important in the identification of cytokines and lymphokine homologs, which are notorious for exhibiting low or undetectable homology at the sequence level (https://www.amazon.com/Homology-Folding-Proteins-Biotechnology-Intelligence/dp/3540636056). For example, members of TNF and TNFR superfamilies, recently identified and characterized in *Drosophila*, are missing in the collation of genes implicated in innate immunity in flies and in mosquito vectors[4][15].

In order to understand the evolution of *Plasmodium* resistance in *An. stephensi*, we have sequenced and assembled the genome of an isofemale line from a strain of intermediate-form from Bangalore (IndInt) displaying diminished susceptibility to *Plasmodium*. We present the assembly of this strain and report an inversion not found in the genomes of the three strains of the type-form[4][5][6][18]. Comparative genomics with other *Anopheles* vectors and deep annotation of genes within this inversion, using fold-prediction algorithms, revealed missed or hidden genes likely to be relevant in *Plasmodium* resistance.

## RESULTS

### Assembly of the genome of IndInt strain

Fourteen generations of Iso-female lines from a parent T6 lab-colony of the intermediate-form of *An. stephensi* (IndInt) were grown, both to achieve homogenity required for high-quality assembly from sequencing of pooled DNA from multiple insects and to enrich for *Plasmodium* resistant phenotype. Multiple tools were used to obtain contig-level assembly of IndInt from 60X coverage of reads sequenced using PacBio Sequel II technologies. The assembly statistics obtained from the various tools are shown in Table 1. In the first round, the best assembly statistics was obtained from WTDBG2 tool with a L50 value of 6 and N50 9.1 Mbp. The next best statistics was shown by the tool Flye assembler with a L50 of 15 and N50 value of 3.9 Mbp.

**Table 1:** Statistics of contig-level assemblies by CANU, FALCON, FLYE, and WTDBG2 tools.

|  | **Assembly**<br>**size**<br>**(Mbp)** | **Total contigs** | **Longest**<br>**contig**<br>**(Mbp)** | **Shortest**<br>**contig**<br>**(bp)** | **Median**<br>**contig**<br>**(Kbp)** | **N50** | **L50** |
| --- | --- | --- | --- | --- | --- | --- | --- |
| **CANU** | 405 | 3464 | 16.6 | 1055 | 44.2 | 48.7 Kbp | 115 |
| **WTDBG** | 205.4 | 789 | 33.08 | 1126 | 15.4 | 9.1 Mbp | 6 |
| **FALCON** | 293 | 851 | 11.4 | 5734 | 87.4 | 1.4 Mbp | 46 |
| **Flye** | 211.7 | 689 | 27.15 | 1007 | 14.6 | 3.9 Mbp | 15 |
| **Flye +**<br>**WTDBG** | 213.3 | 521 | 45.56 | 1018 | 8.3 | 17.04 Mbp | 4 |
| **(Flye +**<br>**WTDBG) +**<br>**SSPACE** | 213.8 | 464 | 55.4 | 1018 | 5.9 | 36.9 Mbp | 3 |

To improve the contiguity of the assembly, the four different contig-level assemblies from CANU, FALCON, FLYE, and WTDBG2[19][20][21][22] were merged in different combinations using the tool Quickmerge[23]. Assembly statistics were computed for each of the merged combinations to determine the best merged assembly (Table 1). During the second round, the assembly with the best statistics was obtained by merging assemblies from FLYE and WTDBG2, with a L50 of 4 and a high N50 of 17 Mbp. The computed genome size of this merged assembly was comparable to the actual size of *An. stephensi* genome, and was used for scaffolding. The scaffolds of the error-corrected contigs were built using the SSPACE assembly tool with simulated mate-pairs of varying sizes from the assembly of IndCh strain as reference. The L50 value for the scaffold level assembly was 3 and the N50 value significantly increased from 17 Mbp to 37 Mbp (Table 1, last row).

In order to check any heterozygosity from potential inversions, haplotype phasing of the assembly of the IndInt strain was done using a set of primary and associated contigs from FALCON and FALCON-unzip tools, respectively[20]. In the presence of heterozygosity, the mapped reads will be split between two contigs that represent the two forms of the heterozygous allele, resulting in reduced coverage. This creates two peaks in the histogram of read counts, one for the haploid level coverage from heterozygous regions and the other for the diploid level coverage from homozygous regions. Fig. 1a with a hump indicates the presence of potential inversion heterozygosity in the genome of IndInt.

**Figure 1.**
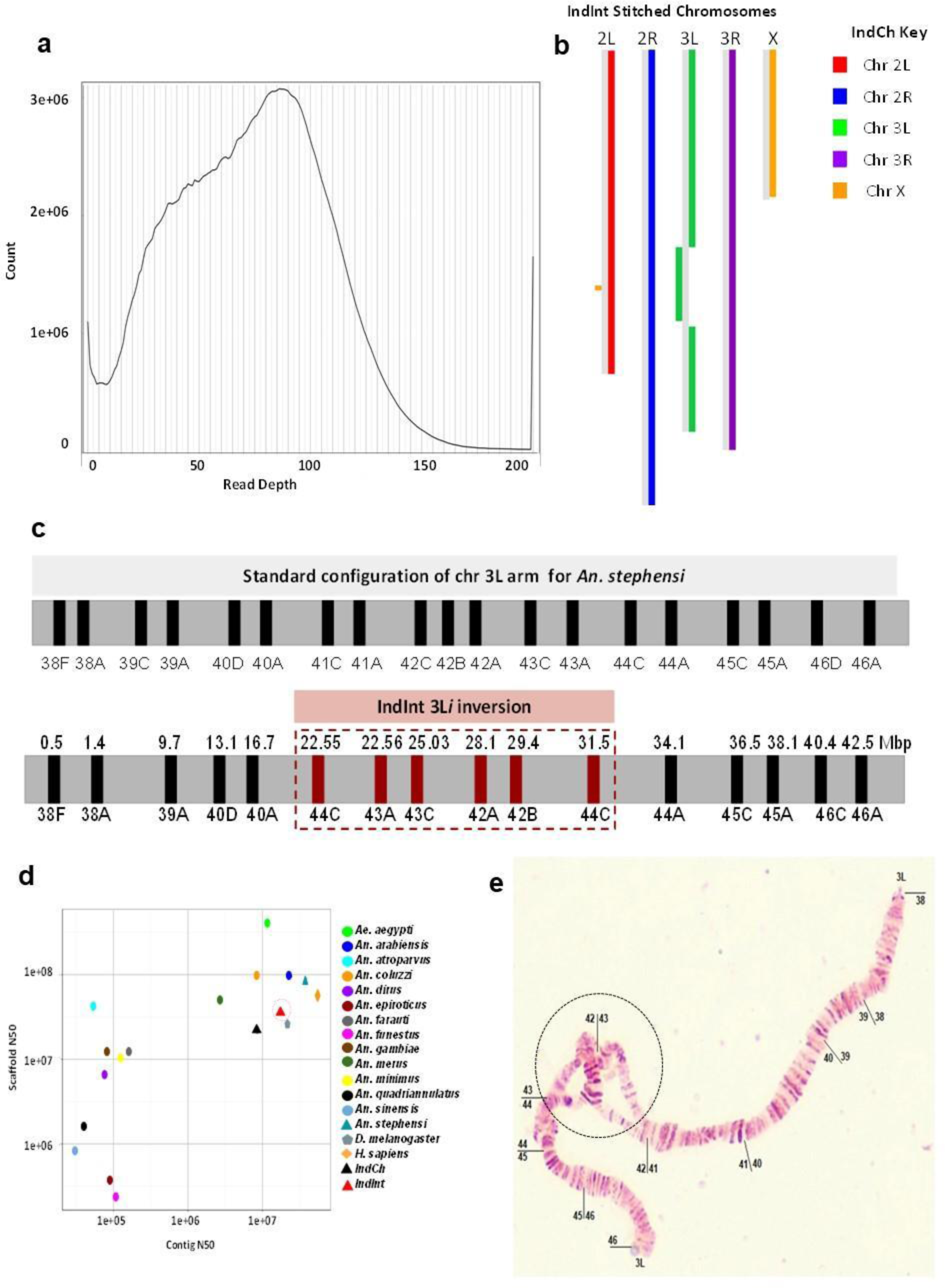
(a) Histogram for the genome assembly of IndInt, showing evidence of heterozygosity by Purge Haplotigs tool. (b) Block diagram of synteny between IndCh and IndInt assembly showing an inversion block in chromosome 3L of IndInt. (c) Top - Orientation of the physical markers as expected in the standard configuration for An. stephensi. Bottom - BLASTN analysis of the assembled IndInt strain revealing the orientation of the physical markers consistent with the 3Li inversion. (d) Quality of IndInt assembly as compared to that of other Anopheles species and completed gold-standard genomes of human and fly. (e) A sample photograph of polytene chromosomes from the IndInt strain showing the loop from the heterozygous inverted configuration of the 3Li.

Synteny between the assemblies of IndInt and IndCh strains is shown in Fig. 1b. The block view clearly suggests that IndInt is homozygous for the standard form of 2R*b* (2R*^+b^*/2R*^+b^*) inversion similar to the IndCh strain. However, in the 3L arm of chromosome 3 IndInt shows a clear presence of an inversion not seen in the IndCh assembly. The locus of the physical markers revealed that the markers from 42A to 44C are inverted in the assembly, which matched with a reported breakpoint for the 3L*i* inversion in *An. stephensi*[8]. Based on the photomaps from several individuals of IndInt strain (see Fig. 1e), it is clear that 48% of the individuals are heterozygous for the 3L*i* inversion (3L*^+i^*/3L*i*), with the rest (52%) being homozygous for the standard form of 3L*i (*3L*^+i^*/3L*^+i^*) (Supplementary Fig. 1). It should be mentioned that the percentage heterozygosity reported here is from an outgrown generation maintained in the lab after the generation used for assembly. Markers 41A, 41B, 41C, and 46D shown in the standard form reported for 3L*i* (top Fig. 1c) are not detectable in the IndInt assembly using the BLASTN tool, perhaps because of limitations of the assembly tools in handling noise generated by reads representing the two arms near the inversion breakpoints from the heterozygous arms.

### Validation of the 2R*b* breakpoint region in IndInt

The IndInt strain is homozygous for the standard form of the 2R*b* inversion, making it the second assembled genome for one of the three possible 2R*b* genotypes, such as *2R^+b^/2R^+b^*. This provides an opportunity to further refine and validate the breakpoint regions described previously by our group[5]. A BLASTN of the sequences from breakpoint regions from IndCh including 1000 bases to the left and right of both breakpoints, against the IndInt assembly, shows 100% identity for all the 2079 bases from the centromere-proximal breakpoint and 99.8% identity for all the 2009 bases from the centromere-distal breakpoint. We also provide the actual alignment with 50+/- nucleotides around the breakpoints in Supplementary Fig. 2. This confirms the 2R*b* genotype to be 2R*^+b^*/2R*^+b^*for IndInt.

### Interrogation of 3L*i* breakpoints in IndInt

In order to define the breakpoints for the 3L*i* inversion, a contact map was generated by mapping the HiC reads from the UCI strain onto the final IndInt assembly. Since UCI is homozygous for the standard (3L*^+i^*/3L*^+i^*) configuration, two expected butterfly-like structures in the chromosome 3L were observed in the contact map (Fig. 2a) defining the location of both the proximal and distal breakpoints of the 3L*i* inversion in the IndInt assembly. By magnifying each of the butterfly structures to a resolution of 5 Kbp per dot, the proximal and distal breakpoints of the 3L*i* inversion were estimated to be between bp 22,545,000 to 22,550,000, and between bp 31,555,000 to 31,560,000 respectively.

**Figure 2:**
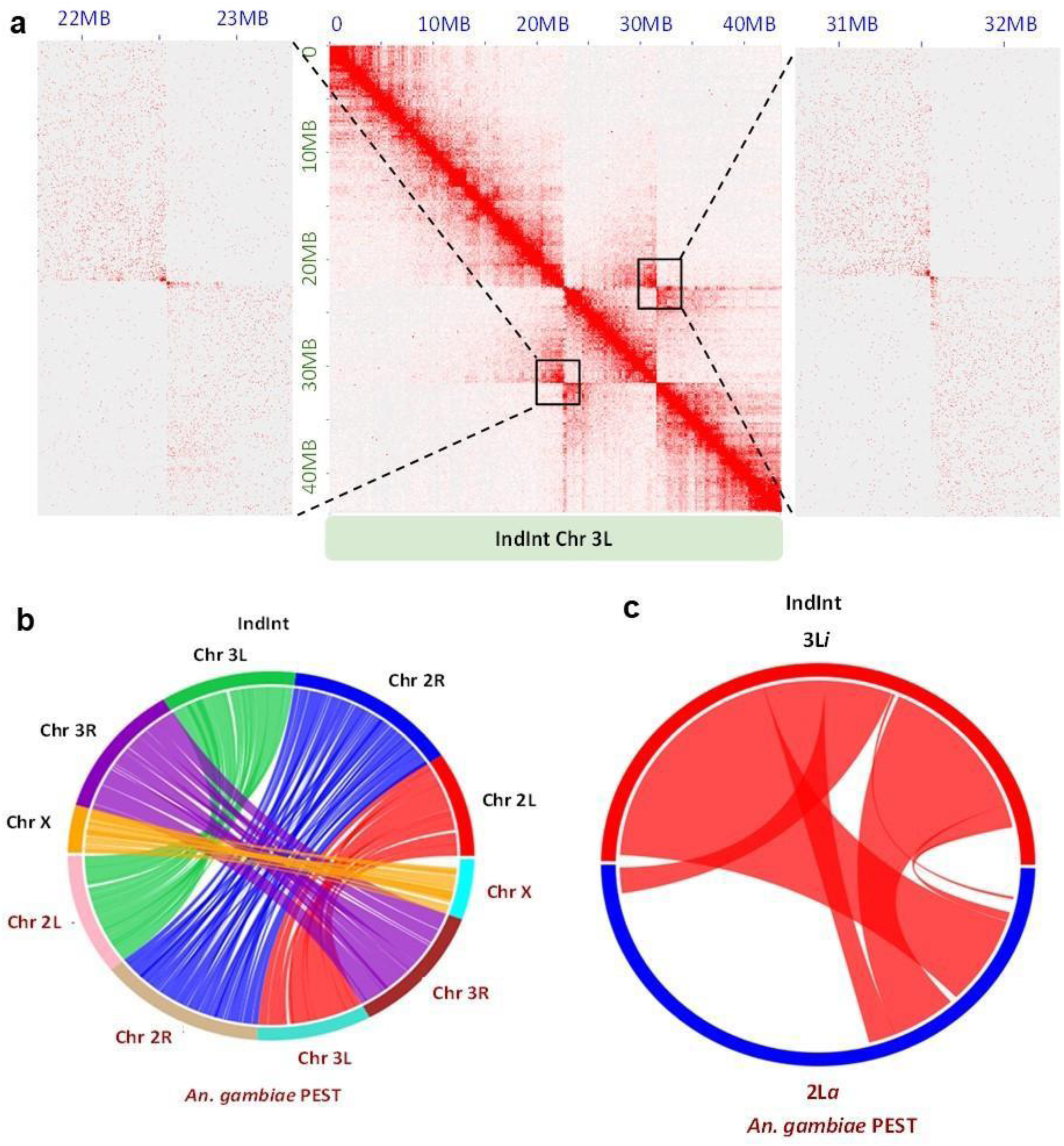
(a) Proximal and distal breakpoints obtained by mapping UCI HiC reads against the chromosome 3L of IndInt. The black boxes represent the centromere-proximal (left) and centromere-distal (right) breakpoints located between the ranges of bp 22,545,000 to 22,550,000 and bp 31,555,000 to 31,560,000, respectively, in the IndInt assembly. Adjacent to the contact map are the enlargements of the butterfly structures, for a resolution of 5 Kbp per dot. (b) Synteny between An. gambiae and IndInt assemblies. (c) Synteny between the An. gambiae 2La and the IndInt 3Li regions.

### Synteny of 3L*i* inversion with 2L*a* of *An. gambiae*

Synteny of the final IndInt assembly was also plotted against the *An. gambiae* PEST strain[24]. Fig. 2b shows that chromosome X, 2R, and 3R of *An. gambiae* is homologous to that of IndInt, respectively. However, the synteny between chromosomal arms 2L and 3L of *An. gambiae* are switched to chromosomal arms 3L and 2L of IndInt, respectively, as reported elsewhere[5]. Fig. 2c shows the synteny between the 2L*a* inversion of *An. gambiae* with the 3L*i* region of IndInt. The 2L*a* region of *An. gambiae* was taken from a previous study[25]. The circle view (Fig. 2c) of this synteny reveals that the entire 3L*i* inversion region of IndInt consisting of 8 Mbp is syntenic to a big part of the 2L*a* inversion region of *An. gambiae* comprising 21 Mbp. Since the 2L*a* inversion of *An. gambiae* is associated with *Plasmodium* sensitivity[11][12], by analogy, one can infer that the 3L*i* of IndInt may also be associated with the *Plasmodium* resistance observed for IndInt strain. Furthermore, it is reported that the strains of *An. gambiae* that are homozygous for the standard form of 2L*a* (2L*^+a^*/2L*^+a^*), show higher vectorial capacity than the corresponding strains that are heterozygous for the inverted form. Similarly, the strain IndCh with higher vectorial capacity is homozygous for the standard form of 3L*i* (3L*^+i^*/3L*^+i^*) and IndInt is heterozygous for the 3L*i* inversion (3L*^+i^*/3L*i*), suggesting the potential role of the 3L*i* inversion in *Plasmodium* sensitivity.

### Functional characterization of genes within the 3L*i* region

Within the 2L*a* region of the *An. gambiae* PEST strain spanning 21 Mbp, there are 1243 genes of which there are 494 genes that are homologous/syntenic to the 3L*i* region of IndInt. Of these, 464 are functionally annotated in IndInt based on sequence level annotation tools, and the remaining 30 genes are uncharacterized due to a lack of sequence homology to known functional genes. Among functionally-characterized genes, we found 36 isoforms of cuticle-forming genes in IndInt. Interestingly, cuticle thickening is associated with pyrethroid & insecticide resistance in other *Anopheles* species[26], while also resisting *Plasmodium* infection[27]. We found 99 missense mutations in the IndInt strain within 23 copies of the cuticle genes, as compared to the IndCh strain of the type form, which has a homozygous configuration for the standard form of 3L*i*, like the UCI strain.

In order to find genes implicated in immune response within the 3L*i* region of IndInt, we intersected the 494 genes within this region with a list of 589 genes reported to be implicated in immunity in insects [15]. Out of these 589 genes, we found only 17 genes within the 3L*i* region (Table 2) including NFkB, a transcription factor implicated in cytokine production in humans.

**Table 2:** Genes within the 3Li region of IndInt that are reported to be implicated in immune response by Waterhouse et al. [15]. Highlighted in cyan is the only gene implicated in TNF-TNFR signaling pathway.

| **IndInt**<br>**Augustus**<br>**ID** | **IndInt**<br>**Chr 3L**<br>**Start** | **IndInt**<br>**Chr 3L**<br>**End** | **Waterhouse et.**<br>**al [15]**<br>**Immune DB ID** | **Function** |
| --- | --- | --- | --- | --- |
| g22033.t1 | 22194979 | 22196095 | AGAP005552 | Peptidoglycan recognition protein |
| g22086.t1 | 22559888 | 22560559 | AGAP007020 | Thioredoxin peroxidase 5 |
| g22107.t1 | 22657471 | 22661294 | AGAP006939 | Armitage protein |
| g22121.t1 | 22702737 | 22707661 | AAEL001675 | Clip domain, serine protease family A |
| g22273.t1 | 24373199 | 24379261 | AGAP006631 | Class A Scavenger Receptor (SRCR domain) with Serine Protease domain |
| g22562.t1 | 26855939 | 26862215 | AGAP006914 | Fibrinogen-related protein 1 |
| g22567.t1 | 26922106 | 26923713 | AGAP006911 | Serine protease inhibitor - Serpin 2 |
| g22568.t1 | 26925252 | 26932268 | AGAP006909 | Serine protease inhibitor - Serpin 1 |
| g22799.t1 | 28434986 | 28435854 | AGAP006790 | Fibrinogen C-terminal domain-containing protein |
| g22840.t1 | 28809510 | 28812635 | AGAP006761 | 3-glucan binding protein |
| g22865.t1 | 28987304 | 28992526 | AGAP006747 | IMD pathway signaling NF-kappaB Relish-like transcription factor |
| g22870.t1 | 29022137 | 29026993 | AGAP006743 | Unspecified product<br>BLASTP : Similar to ficolin 1 |
| g22921.t1 | 29370252 | 29370571 | AGAP006722 | Unspecified product<br>BLASTP : Similar to Cecropin B1 |
| g23080.t1 | 30464940 | 30470336 | AGAP005672 | Staphylococcal nuclease domain-containing protein 1 |
| g23109.t1 | 30680813 | 30689983 | AGAP005652 | ATP-dependent RNA helicase DDX5/DBP2 |
| g23151.t1 | 30998165 | 31003426 | AGAP005625 | Class A Scavenger Receptor (SRCR domain) with Serine Protease domain |
| g23220.t1 | 31575970 | 31576777 | CPIJ004568 | Similar to Cu-chaperone |

The genes implicated in *Plasmodium* resistance in *An. gambiae*[28] LRIM1, APL1C, and TEP1 are not within the 3L*i* region in IndInt. The TEP1 protein in IndInt is located on chromosomal arm 2L. The LRIM1 gene is outside the 3L*i* region, while the gene APL1C is located immediately outside the distal breakpoint of the 3L*i* region. The LRIM1 gene has leucine-rich repeats and forms a disulfide-bonded complex with APL1C (PDB ID: 3OJA [28]), which stabilizes the mature TEP1 and promotes its binding to parasites to trigger pathogen destruction. Also, more recently, a knockout of LRIM1 in *An. stephensi* was shown to play a role in vector competence[29]. Furthermore, LRIM1 and APL1C are within the 2L*a* region in *An. gambiae,* which is three times larger than the 3L*i* region in IndInt. In IndInt, the LRIM1 has four and six missense mutations compared to IndCh and UCI strains of type-form, respectively. However, since the gene APL1C shows no missense mutation in IndInt compared to IndCh, we expected other unannotated paralogs of LRIM1 within the 3L*i* region of IndInt. Deep annotation of the 30 functionally uncharacterized genes within the 3L*i* region shown in Supplementary Fig. 3 with fold-prediction algorithms, revealed two novel leucine-rich proteins, g22432 and g22212, similar in structure to LRIM1 (Supplementary Fig. 3a and 3b). These two LRIM1-like genes could potentially work with the intact APL1C gene in IndInt to launch a host defense system to antagonize the malaria parasite. The high conservation of these, otherwise uncharacterized genes across *Anopheles* species, shown in the multiple sequence alignments of g22432 and g22212, suggests functional conservation across species (Supplementary Fig. 4a and 4b).

Recently, members of LPS-induced TNF-alpha factors, LITAFs, have been shown to regulate anti-*Plasmodium* immunity in *An. gambiae*[30][31]. A search in the assembled genome of *An. stephensi* identified four members of LITAF-like transmembrane genes (Supplementary Text 1). LITAFs are known to send signals to the nucleus to transcribe TNF-alpha molecules in humans [32]. In fact, LPS-induced animal models were used in developing drugs against TNF-alpha to treat auto-immune disorders. We were also able to find homologs of all the genes involved in this signaling pathway in IndInt (Supplementary Text 2).

We found an eiger homolog, g22826, which is within the 3L*i* region of IndInt and is missing in the reported list of 361 genes with immune repertoire in *An. stephensi* [4]. A crystal structure of the complex of eiger and its receptor from *Drosophila* has revealed that eiger belongs to the TNF superfamily and its two receptors, wengen and grnd, are members of the TNFR superfamily(6ZT0 [33]). A search for an active TNFR-like gene in *An. stephensi,* using the HMM model (PF00020) of the cysteine-rich domain (CRD) from PFAM database, identified two potential hits g1129 and g18030 in IndInt. It should be mentioned that while wengen displays detectable homology with TNFRSF, predicted grnd homologs from many sequenced species, including malaria vectors, remain uncharacterized by sequence-based annotation tools. The gene, g1129 from chromosome X of the IndInt strain encodes a wengen homolog of *Drosophila*, with a well-defined extracellular CRD, and a transmembrane region. The grnd homolog (g18030) in the IndInt strain consists of an intact CRD region sandwiched between a predicted signal sequence and transmembrane region (Supplementary Fig 5c and 5d). We found that the eiger gene has a frame-shift mutation at the tail-end of the very conserved folding domain in the IndCh assembly. Also, in the UCI and STE2 strains, the predicted eiger homologs are missing the critical N-terminal transmembrane region (Supplementary Fig. 6). In Fig. 3a, a 3D model of the eiger gene from IndInt, built using RoseTTaFold tool[16], is shown superimposed on the crystal structure of human TNF-TNFR complex (PDB: 1TNR,[34]) on the left, along with a wengen model built using homology to the second domain of TNFR-1A. Similarly, the eiger model is shown superimposed on the crystal structure of the eiger-grnd complex (PDB: 6ZT0 [33]), along with a model of grnd built using homology to grnd of *Drosophila* in Fig. 3b. The choice of templates for comparative modeling of the CRDs of the two receptors is based on the conserved cysteine patterns of the ligand binding regions, which are different for the two receptors wengen and grnd. Interestingly, the wengen binding domain has a cysteine pattern similar to TNFR-1A of humans and that of grnd is similar to that from TNFR-1B, suggesting an evolutionary-conserved link between eiger and TNF-alpha mediated signaling in arthropods and vertebrates respectively.

**Figure 3:**
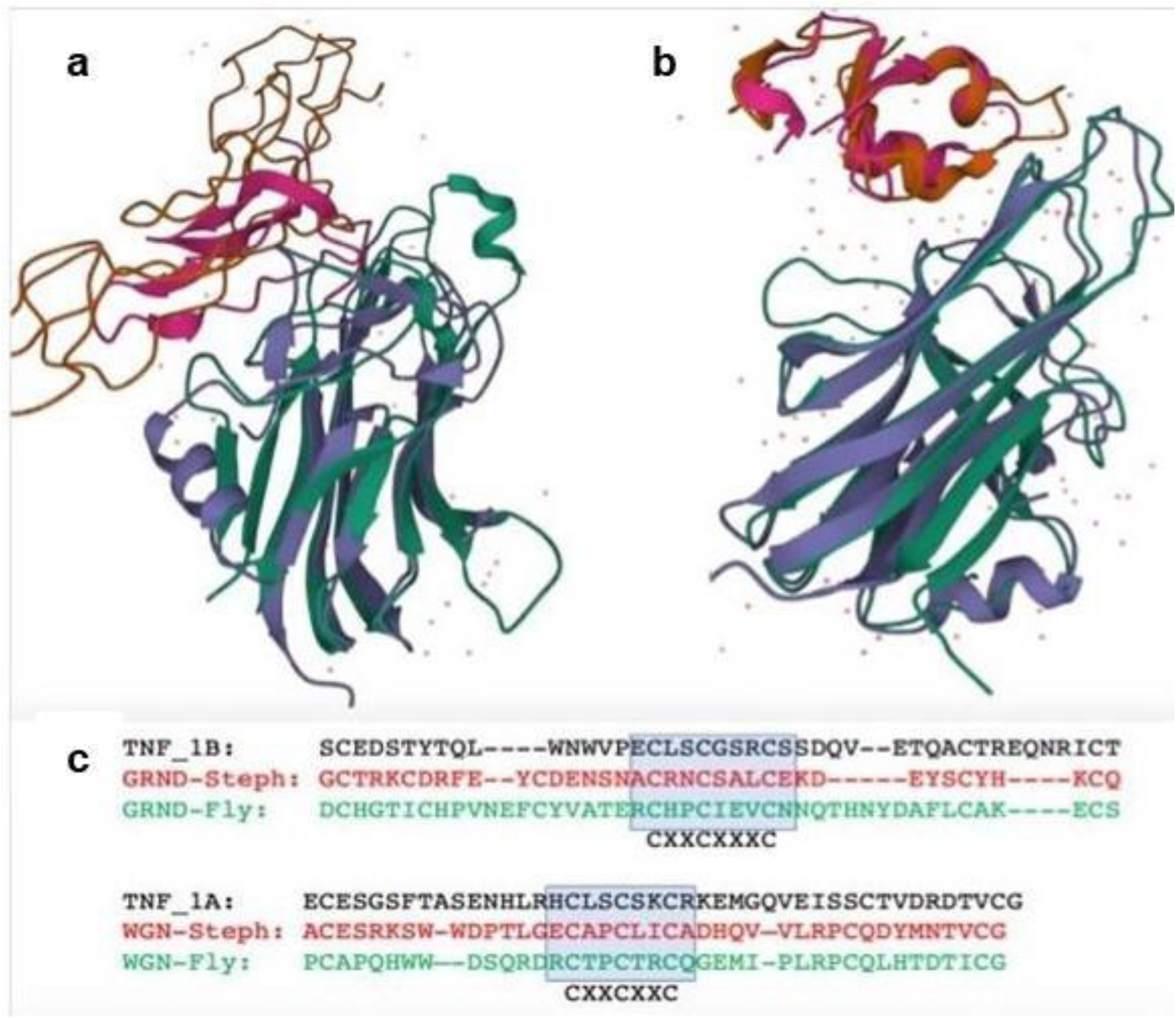
(a) De novo model of eiger and comparative model of wengen superimposed on the crystal structure of the complex of TNFb-TNFR-1A (1tnr) (b) De novo model eiger and comparative model of grnd superimposed on the crystal structure of the complex of eiger and grnd from Drosophila (6zt0) (c) Sequence alignment of wengen and grnd against TNFR-1A, -1B and grnd from Drosophila used in comparative modeling.

### A functional TNF-alpha/TNFR pathway in malaria vectors with lower vectorial capacity

It is now well established that the *Drosophila* genome harbors two functional TNFR receptors and a single ligand, eiger, belonging to the TNF-TNFR system[33]. In order to predict whether the homologs of the eiger and its cognate receptor genes in *An. stephensi* (IndInt) are potentially functional, we used signal peptide and transmembrane predictions on the respective protein sequences derived from IndInt (Supplementary Fig. 5). The eiger homolog shows a transmembrane region at its N-terminus (Supplementary Fig. 5a) consistent with type II configuration common to all members of the TNF superfamily across organisms. The two receptor homologs show clear transmembrane regions (Supplementary Fig. 5b and 5c) following the cysteine-rich folding domain. However, only grnd, but not wengen, has a predicted signal peptide signature (Supplementary Fig. 5d) N-terminal to the CRD, consistent with the localization of these two genes in *Drosophila,* where wengen is almost always localized in intracellular vesicles[33]. We also found the homolog of the TACE gene in IndInt, which is a TNF-alpha processing metalloprotease that converts the type-II membrane-anchored ligand into its soluble form in humans [35]. In the eiger gene from IndInt, we also found a cleavage site (AQ) N-terminal to the folding domain that is known to be specific to TACE [36].

In order to show that all the genes in the cytoplasm necessary for eiger-wgn-grnd mediated signaling are encoded by the genome of IndInt, we identified homologs of all the pathway genes from TNF-TNFR signaling pathway from KEGG (Supplementary Text 2). Fig. 4 shows genes implicated in wengen-based (left) and grnd-based (right, hypothesis) based on the homology of the receptors to TNFR-1A and -1B, respectively. Several genes are disrupted in the other strains of type-form viz IndCh, UCI and STE2, as shown in Fig. 4, Supplementary table 1 and Supplementray table 2. The homologs for genes marked by red and purple stars are disrupted and absent, respectively, in the IndCh genome, potentially disrupting eiger-mediated signaling pathway. Genes with blue stars are missing in all 4 assembled genomes, suggesting that these pathways may have evolved in vertebrates after the invertebrate-vertebrate split.

**Figure 4:**
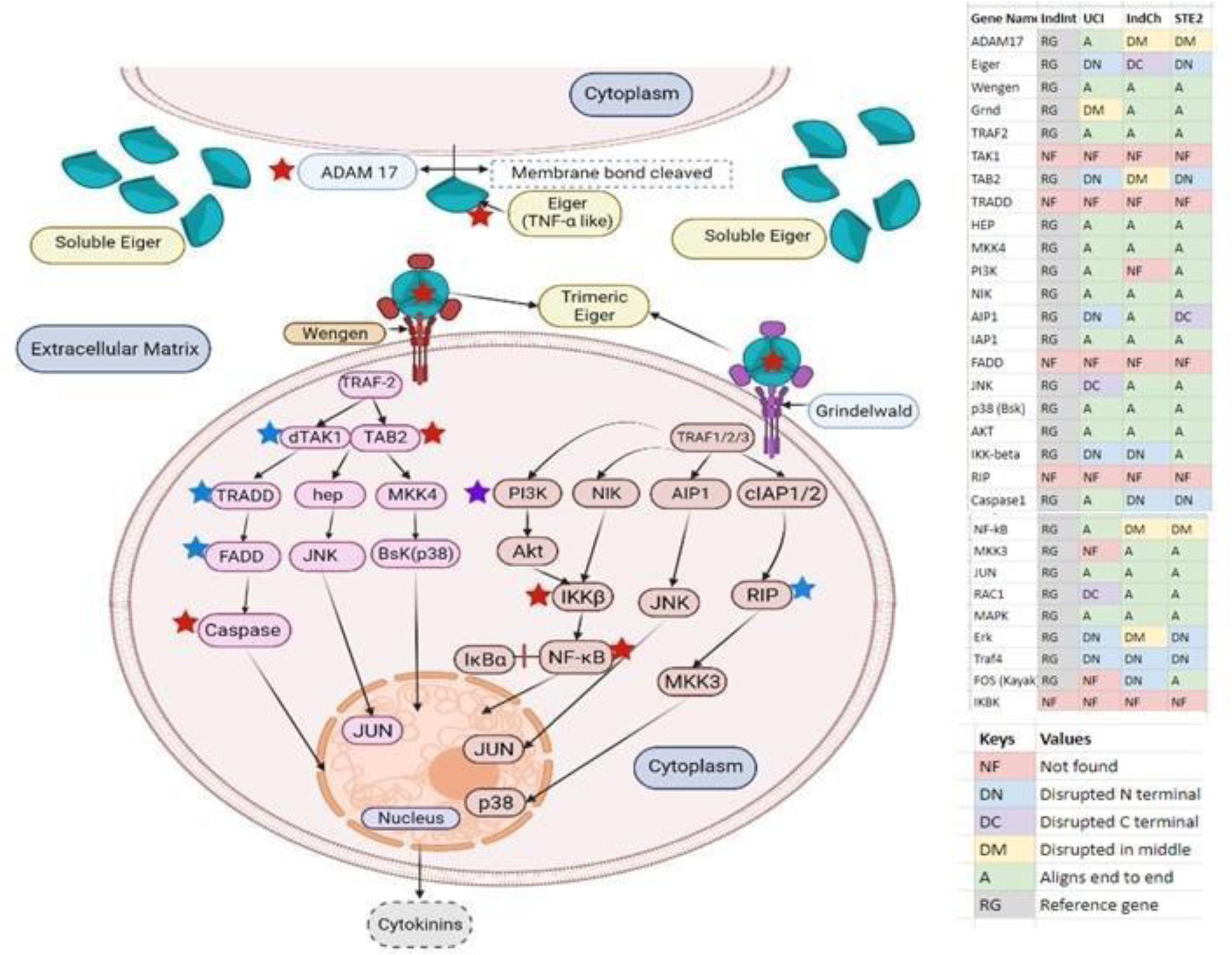
Pathways leading to eiger-grnd/wgn-mediated signaling collated from TNF-TNFR and eiger-wgn-mediated signaling pathways obtained from the KEGG database (https://www.genome.jp/pathway/hsa04668) and validated using other sources [37][38]. Red stars represent the disrupted genes in the IndCh genome, purple star is the gene missing in IndCh but present in IndInt, blue stars are the missing pathway genes in all four assembled An. stephensi genomes. The inset table shows the status of the genes in all 4 strains (IndInt, IndCh, UCI and STE2) of An. stephensi for which a high-quality genome is available. (Template for designing the pathway diagram was adapted from https://app.biorender.com/biorender-templates)

It should be mentioned that a majority of these genes including eiger, grnd, and wengen are missing in the collated list of 589 immune related genes in insects[15] and, more specifically, missing from a list of 361 immune related genes in *An. stephensi* reported recently in the UCI strain[4].

## DISCUSSION

We report a high-quality genome assembly of an isofemale line of the intermediate-form of *An. stephensi*, IndInt, displaying low vectorial capacity relative to the strain IndCh of type-form (manuscript under preparation). A comparative analysis of this assembly with those of type-forms, displaying higher vectorial capacity, reveals a 3L*i* inversion (in a heterozygous configuration) that is uniquely present only in the IndInt strain. As shown in Fig. 2c, this inversion is syntenic to the 2L*a* inversion region in *An. gambiae*, which is shown to be associated with *Plasmodium* sensitivity[11]; thus providing an opportunity to identify intact genes that may be offering resistance to *Plasmodium*. Here, for the first time, we report an intact TNF-TNFR system in malaria vectors with eiger, a member of the TNF superfamily, present within this inversion region (Figure 3b). Considering that the eiger gene in the IndCh strain harbors a frameshift mutation within the structurally folding domain, it is tempting to predict a role for this gene in the innate immune response of *An. stephensi* against *Plasmodium* in IndInt strain. Similarly, the predicted eiger gene in STE2 and UCI strains are both disrupted in the N-terminal lacking transmembrane region and several genes in the eiger pathway are disrupted in IndCh strain (Fig 4, Supplementary table 2). These observations suggest a testable correlation between high vectorial capacity in *An. stephensi* strains with a dysfunctional eiger-mediated signaling pathway, which has a lower vectorial capacity for *Plasmodium* with a potentially functional eiger-signaling pathway. Furthermore, the increased expression of eiger-wgn genes in response to infection by Nora virus in *Drosophila*[39], demands an exploration of the role of this complex in *Plasmodium* resistance in *An. stephensi* and in other malaria vectors, in general.

The gene TNF-alpha and its two cognate receptors in humans, TNFR-1A and -1B, are implicated both in adaptive and innate immune response. Indeed, drugs inhibiting this pathway are now available to treat various autoimmune diseases such as rheumatoid arthritis, Crohn’s disease, and psoriasis. The two cognate receptors of eiger, wengen and grnd, show homology in their cysteine patterns to TNFR-1A and -1B respectively (Fig. 3c), suggesting evolutionary conservation of this signaling pathway. Consistent with the ability of the eiger gene to participate in innate immune response analogous to signaling by TNF-alpha, we found genes in *An. stephensi* (IndInt) that are homologous to genes implicated in TNF-alpha mediated signaling (Fig 4). We have found homologs of just about every gene required for eiger-based signaling similar to TNF-alpha in the IndInt assembly of *An. stephensi* (Fig. 4). Considering that many genes in the TNF-alpha pathway are involved in multiple processes, it is surprising that a majority of these are missing from the list of genes with immune repertoire reported for insects [15] (Supplementray Table 1).

It is known that human TNF-alpha reduces *Plasmodium falciparum* growth and activates calcium channels in human malarial parasites[40]. It is reasonable to hypothesize that mosquito vectors may also elicit similar responses via the eiger-wengen-grnd signaling pathway in parasites to control their growth. It is well established that Ca^+2^ mediated signaling is correlated with the developmental stages of the parasite life cycle[41]. The eiger and its receptor homologs in *Drosophila* uses JNK pathway in a manner similar to human TNF-alpha. For example, in *Drosophila,* a grnd null mutant completely abolishes cell-death[33]. Furthermore, the eiger mediated cell-death in flies could be reversed by JNK inhibitors[33]. These observations support the hypothesis that a functional eiger pathway in malaria vectors may reduce the growth of *Plasmodium falciparum,* similar to the action of TNF-alpha in the human host.

*Drosophila* and mosquitoes diverged some 350 million years ago, suggesting that the TNF-TNFR system-based innate immunity may have evolved before this point of inflection. A search for TNF- and TNFR-like molecules in the Alpha-fold database of *C. elegans* reveals ∼14 TNF-like molecules without any TNFR-like homologs. The TNF-like molecules in *C. elegans* are actually cerebellin homologs. Interestingly, cerebellins are also both type 2-membrane proteins and form 3 fold functional oligomers just like members of the TNF superfamily. As proof of this, there are homologs of glutamate receptors and neurexin genes encoded by *C. elegans*, which are involved in forming synaptic structures in rats [42]. Since the TNF-TNFR system is conserved in all insects and vertebrates, missing/lost only in *nematodes*, innate immunity by the TNF-TNFR system may predate the invertebrate-vertebrate split and TNF-like fold evolution may even be older.

## CONCLUSION

The three forms of *An. stephensi* (type, intermediate and mysorensis), displaying varying vector competence, which are clearly identifiable by their number of egg ridges, offer an ideal system to discover genes implicated in *Plasmodium* resistance by interrogating genomes of strains with low vectorial capacity. Here, by assembling the genome of an intermediate-form (IndInt) and comparing it to that of a type-form (IndCh), we report a unique 3L*i* inversion in IndInt, which is syntenic to the 2L*a* inversion in *An. gambiae* previously implicated in *Plasmodium* resistance. This has allowed, for the first time, identification of TNF-alpha homolog, eiger, in malaria vectors along with homologs of genes required for downstream-signaling to offer an innate immune response. Most strikingly, IndCh strain, with high vectorial capacity for *Plasmodium*, harbors disruptive mutations in one or more genes of the eiger-grnd-wengen pathway. These observations strongly suggest that disruption of eiger-based signaling in the type-forms of *An. stephensi* may account for their high vectorial capacity predicated upon a reduction in innate immunity to combat *Plasmodium* infections.

## MATERIALS AND METHODS

### Maintenance of *An. stephensi* in insectary

Larvae of *An. stephensi* were collected from Sriramanahalli area of Bangalore rural (13.4310°N, 77.3310°E) from the state of Karnataka, India, and successfully colonized in the insectary. Larvae were provided with larval food prepared by mixing Brewer’s yeast and dog biscuits at a ratio of 30:70. Pupae were bleached with 1% sodium hypochlorite for 1 min and kept in adult rearing cages (Bugdorm-4S3030) (W32.5 × D32.5 × H32.5 cm). The adults were fed on a mixture of 8% sucrose, 2% glucose solution mixed with 3% multivitamin syrup (Polybion LC®)[43]. Adults were maintained in the insectary at 28±1°C, RH 75±5% and 12:12-h day and night photoperiod cycles. Adult females were provided with blood on day 7 or 8 post-emergence to obtain eggs for continuing next progeny. Eggs were collected on day 4 in an ovicup containing water and lined with filter paper at the inner margin. Larvae were hatched out after 36 to 48 h and transferred to the rearing trays (L39 x B30 cm; Polylab, Catalog no. 81702) with larval food.

### Establishment of IndInt iso-female line of *An. stephensi*

About 10-15 gravid females were separated from the intermediate line[44]. On day 3, each gravid female was transferred carefully to a single ovicup. Each ovicup was covered with a nylon net. A cotton ball soaked with sugar solution was kept on the top of each ovicup. The line from single female (G_0_) was selected to generate the iso-female lines that laid the highest number of eggs and had highest percent hatchability with egg ridge numbers 15-17. The eggs thus collected from the single female were allowed to hatch, and the larvae were provided with larval food and reared to adults as per the protocol mentioned earlier. Emerged adult siblings (G_1_) were kept in a mosquito rearing cage and allowed to inbreed (sibling mating). Mated females were blood fed, and the same procedure was followed as mentioned earlier and continued till further generations. As of July 2022, the IndInt iso-female colony has been maintained in our insectary for 55 generations.

### Karyotyping of chromosome 3 of IndInt strain

Polytene chromosomes were prepared from the ovarian nurse cells collected from semi-gravid females of the IndInt iso-female lines as per the method of Ghosh & Shetty, which was adapted from our recent publication[5]. The semi-gravid females were anesthetized and placed on a microslide in a drop of phosphate buffer. The ovaries were pulled out gently and fixed in modified Carnoy’s fixative (methanol: acetic acid, 3:1) for 2-3 min. The remaining mosquito body was preserved in an Eppendorf tube with a few drops of phosphate buffer solution and kept at - 20°C for PCR analysis. After fixation, the ovaries were stained with lacto acetic orcein for 15-20 min. After staining, 60% acetic acid was added to it and a clean coverslip was placed on top of the stained sample. Gentle pressure was applied on the cover glass for squashing. The edges of the coverslip were sealed with nail polish to avoid evaporation. The slides were examined under a microscope using a 40X objective lens for chromosome inversions. The inversion nomenclature and their frequency were recorded[45].

### Assembly

PacBio reads from the IndInt strain were assembled independently using FLYE[21] and WTDBG2[22] assemblers. The resulting assemblies were combined using Quickmerge[23] where the FLYE assembly was taken as the query and the WTDBG2 assembly served as the reference. Two rounds of Arrow polishing (PacBio gcpp v2.0.2) using PacBio reads, followed by two rounds of Pilon polishing (v1.22) using 100x Illumina reads, were carried out on the merged assembly. In order to proceed with reference based de novo assembly, simulated mate-pair reads were generated from the IndCh standard assembly (DOI: 10.1038/s41598-022-07462-3) using the tool WGSIM (https://github.com/lh3/wgsim) for the sizes 50 Kbp, 100 Kbp, 250 Kbp, 500 Kbp, 1 Mbp, 2.5 Mbp and 5 Mbp. These simulated mate-pair reads along with the contig level assembly from FLYE+WTDBG2 served as input to obtain a scaffold level assembly using the SSPACE[46] tool. DNA physical marker sequences for each chromosome arm published by Jiang *et al*[18]. were downloaded and aligned against the scaffolds using BLAST, in order to assign the chromosomal positions to the scaffolds. The scaffolds were stitched based on the order and orientation of the markers into pseudo-chromosomes. 100 N’s were added between two scaffolds during stitching. Physical marker sequences were realigned to the stitched chromosomes for validation.

### Haplotype phasing

The assemblers FALCON and FALCON-Unzip[20] were used to phase the IndInt genome into haplotypes. Raw IndInt PacBio reads were used by the FALCON assembler to produce a set of primary contig files and an associate contig file representing the divergent allelic variants. The output of primary and associate contig files were then used by FALCON-Unzip to produce partially phased primary contigs (all_p_ctg.fa) and fully phased haplotigs (all_h_ctg.fa), which represent divergent haplotypes. Polishing was performed on the phased contigs by FALCON-Unzip using Arrow. This procedure was adapted from our recent publication[5].

### Purge haplotigs

The tool ‘Purge Haplotigs’[47] was used to determine the degree of heterozygosity in the IndInt strain. PacBio reads of IndInt strain were mapped to the consensus primary assembly (cns_p_ctg.fasta) obtained from FALCON-Unzip to obtain an aligned BAM file. Coverage analysis was performed on the BAM file to generate a read-depth histogram.

### Identification of 3L*i* breakpoints in IndInt

The tool HiC-Pro[48] was used to generate a contact map by mapping the HiC reads from the UCI strain onto the genome of IndInt with different inversion genotype to produce a butterfly-like structure representing the breakpoints, which were visualized and extrapolated from the contact map of the tool Juicebox[49]. The breakpoints were validated by mapping the PacBio and Illumina reads onto the respective genomes. Alignment using ‘unimap’ was performed between the genomes of the IndInt and UCI strains, to determine the precise breakpoints for the 3L*i* inversion to consolidate with the findings from HiC-Pro.

### Gene annotation

AUGUSTUS (version 3.2.3)[50], an eukaryotic gene prediction tool, was used to find protein-coding genes for all the assembled genomes. The model organism closest to *An. stephensi, Ae. aegypti,* which was made available by AUGUSTUS, was used for gene prediction. This procedure was adapted from our recent publication[5].

### Structural prediction of genes of unknown functions

The genes obtained from AUGUSTUS were used as an input for RoseTTAFold (https://robetta.bakerlab.org/, version August 2021), a deep learning-based protein modeling method. High confidence models obtained from Robetta were used as an input for PyMOL (Version 2.0 Schrödinger, LLC). Regions conserved across the models were identified after structural alignment in Pymol as putative protein domains. Protein contact maps were generated using BIOVIA Discovery Studio Visualizer (v21.1.0.20298) to visually confirm the conserved regions and check for short and long-range interactions in the identified domains. The identified domains were given as input in both sequence and structure-based comparison softwares. Structural comparison was done using DALI (DaliLite.v5) and sequence comparison was done using online utilities such as SwissModel Expasy, Conserved Domain Database from NCBI, InterPro, HMMER Homology Search, and Pfam Sequence Search. Default parameters were used for all the tools. The DALI result showing optimal RMSD value, and seen as hits in the sequence-based tools as well, was chosen as the possible domain.

## DATA AVAILABILITY

The raw PacBio/Illumina reads used in the assembly and the IndInt chromosome level assembly are uploaded onto the NCBI server under the BioProject ID: PRJNA757559.

## AUTHOR CONTRIBUTION

SSri for overseeing bioinformatics work, setting the tone and writing of the manuscript

SD for writing the first draft, taming the assembly and discovery of 3L*i* inversion

CG for creating the IndInt isofemale line and generating photomaps SSingh for contig level assemblies using various methods

AT for characterizing candidate genes

AG characterizing breakpoints and TNF-TNFR pathway work

HM, AJ, MK, AC, RB, VS for deep annotation of 3L*i* region using RoseTTaFold prediction

SM Insectary work

NK, SK insectary work and for vectorial capacity in both standard and heterozygous 3L*i* inverted individuals

SSwa for overseeing the entire insect work

SS for conceptualization and contributions to the manuscript

## ACKNOWLEDGEMENT

The authors thank Nucleome Informatics Pvt. Ltd. for generating Pacbio reads and Illumina reads for the IndInt strain. The authors thank Tata Institute for Genetics and Society (TIGS) India for funding researchers involved in this work and for sequencing and the Government of Karnataka for funding the computational infrastructure at IBAB.

