## Supplementary Figures and Text for "Identification of a TNF-TNFR-like system in malaria vectors (*Anopheles stephensi*) likely to influence *Plasmodium* resistance"

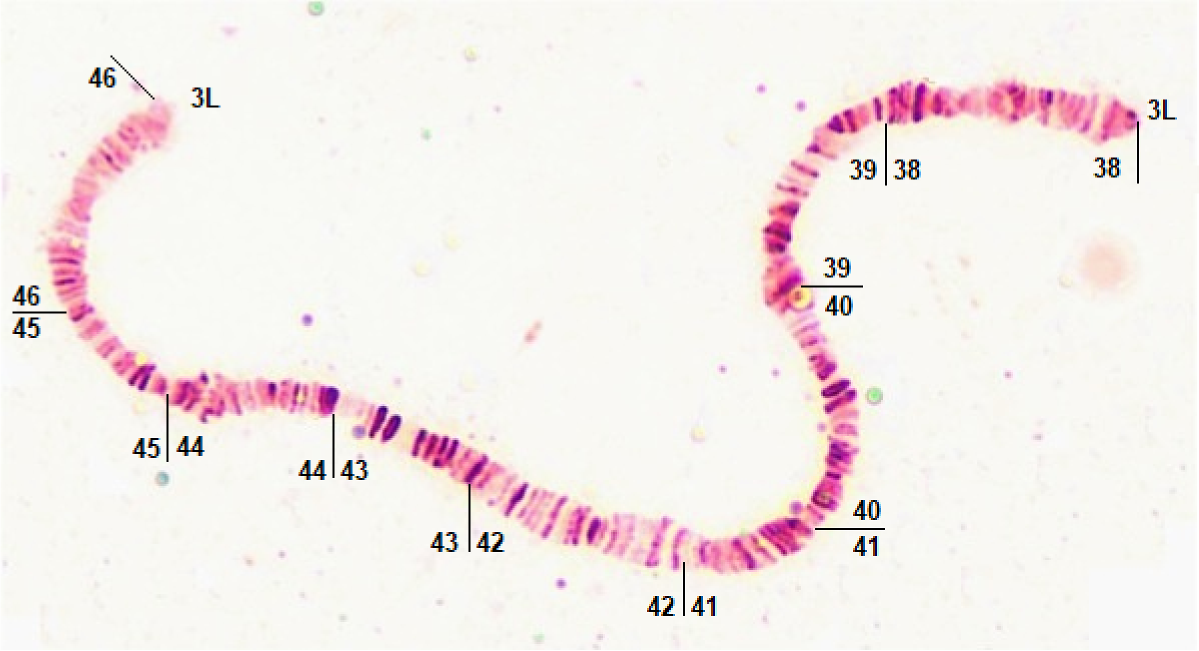


*Supplementary figure 1 : Photomap showing the standard form of the 3Li region in the IndInt strain, supporting 52% homozygous for the standard form of 3Li.*

**78 bp :**

```
Database: IndInt_chr2R.fasta
           1 sequences; 55,427,671 total letters


Query= IndCh_Astel_HiC_Chr2R

Length=55103326
                                                                      Score        E
Sequences producing significant alignments:                          (Bits)     Value

chr2R                                                                 331        1e-90


>chr2R
Length=55427671

 Score = 331 bits (179),  Expect = 1e-90
 Identities = 179/179 (100%), Gaps = 0/179 (0%)
 Strand=Plus/Minus

Query  38082990  GGCAGAAGTGAGTGAATGACTTAAATCTTTGAAATGAATTAGATGATTTAAATCATCACG  38083049
                 ||||||||||||||||||||||||||||||||||||||||||||||||||||||||||||
Sbjct  17244809  GGCAGAAGTGAGTGAATGACTTAAATCTTTGAAATGAATTAGATGATTTAAATCATCACG  17244750

Query  38083050  CAGCGTTATTTCAGCTCACTTTTGATTTAATTCGCCATGCCGATTAACCACTCACTCACT  38083109
                 ||||||||||||||||||||||||||||||||||||||||||||||||||||||||||||
Sbjct  17244749  CAGCGTTATTTCAGCTCACTTTTGATTTAATTCGCCATGCCGATTAACCACTCACTCACT  17244690

Query  38083110  ACTTTTTAACTCACTCACTATGAATGGCTCTCTCGTGAATTAAACGAGTGAGTTAAAGG  38083168
                 |||||||||||||||||||||||||||||||||||||||||||||||||||||||||||
Sbjct  17244689  ACTTTTTAACTCACTCACTATGAATGGCTCTCTCGTGAATTAAACGAGTGAGTTAAAGG  17244631
```

### 78 bp Reverse Complement :

```
Database: indint_rev_chr2R.fasta
           1 sequences; 55,427,671 total letters


Query= IndCh_Astel_HiC_Chr2R

Length=55103326
                                                                      Score     E
Sequences producing significant alignments:                          (Bits)  Value

chr2R                                                                 331     1e-90


>chr2R
Length=55427671

 Score = 331 bits (179),  Expect = 1e-90
 Identities = 179/179 (100%), Gaps = 0/179 (0%)
 Strand=Plus/Plus

Query  38082990  GGCAGAAGTGAGTGAATGACTTAAATCTTTGAAATGAATTAGATGATTTAAATCATCACG  38083049
                 ||||||||||||||||||||||||||||||||||||||||||||||||||||||||||||
Sbjct  38182863  GGCAGAAGTGAGTGAATGACTTAAATCTTTGAAATGAATTAGATGATTTAAATCATCACG  38182922

Query  38083050  CAGCGTTATTTCAGCTCACTTTTGATTTAATTCGCCATGCCGATTAACCACTCACTCACT  38083109
                 ||||||||||||||||||||||||||||||||||||||||||||||||||||||||||||
Sbjct  38182923  CAGCGTTATTTCAGCTCACTTTTGATTTAATTCGCCATGCCGATTAACCACTCACTCACT  38182982

Query  38083110  ACTTTTTAACTCACTCACTATGAATGGCTCTCTCGTGAATTAAACGAGTGAGTTAAAGG  38083168
                 |||||||||||||||||||||||||||||||||||||||||||||||||||||||||||
Sbjct  38182983  ACTTTTTAACTCACTCACTATGAATGGCTCTCTCGTGAATTAAACGAGTGAGTTAAAGG  38183041
```

## 8 bp :

```
Database: IndInt_chr2R.fasta
           1 sequences; 55,427,671 total letters


Query= IndCh_Astel_HiC_Chr2R

Length=55103326
                                                                      Score     E
Sequences producing significant alignments:                          (Bits)  Value

chr2R                                                                 202     6e-52


>chr2R
Length=55427671

 Score = 202 bits (109),  Expect = 6e-52
 Identities = 109/109 (100%), Gaps = 0/109 (0%)
 Strand=Plus/Minus

Query  21514292  ATTTAAGGTTGACCAATTCCTGAGAAACACGGGTAAGAATATAATCCAATCATATCTTTA  21514351
                 ||||||||||||||||||||||||||||||||||||||||||||||||||||||||||||
Sbjct  33742489  ATTTAAGGTTGACCAATTCCTGAGAAACACGGGTAAGAATATAATCCAATCATATCTTTA  33742430

Query  21514352  GCGAGCTGCAAATTAGTGATTTTCCTAGAACACTGGCAACACCGCCAAC  21514400
                 |||||||||||||||||||||||||||||||||||||||||||||||||
Sbjct  33742429  GCGAGCTGCAAATTAGTGATTTTCCTAGAACACTGGCAACACCGCCAAC  33742381
```

**8bp Reverse Complement :**

```
Database: indint_rev_chr2R.fasta
           1 sequences; 55,427,671 total letters


Query= IndCh_Astel_HiC_Chr2R

Length=55103326
                                                                      Score     E
Sequences producing significant alignments:                          (Bits)  Value

chr2R                                                                  202     6e-52


>chr2R
Length=55427671

 Score = 202 bits (109),  Expect = 6e-52
 Identities = 109/109 (100%), Gaps = 0/109 (0%)
 Strand=Plus/Plus

Query  21514292  ATTTAAGGTTGACCAATTCCTGAGAAACACGGGTAAGAATATAATCCAATCATATCTTTA  21514351
                 ||||||||||||||||||||||||||||||||||||||||||||||||||||||||||||
Sbjct  21685183  ATTTAAGGTTGACCAATTCCTGAGAAACACGGGTAAGAATATAATCCAATCATATCTTTA  21685242

Query  21514352  GCGAGCTGCAAATTAGTGATTTTCCTAGAACACTGGCAACACCGCCAAC  21514400
                 |||||||||||||||||||||||||||||||||||||||||||||||||
Sbjct  21685243  GCGAGCTGCAAATTAGTGATTTTCCTAGAACACTGGCAACACCGCCAAC  21685291
```

*Supplementary figure 2 : Pairwise alignment using BLASTN showing the validation of 2Rb breakpoints between the IndCh and IndInt strains.*

**g22432.t1**

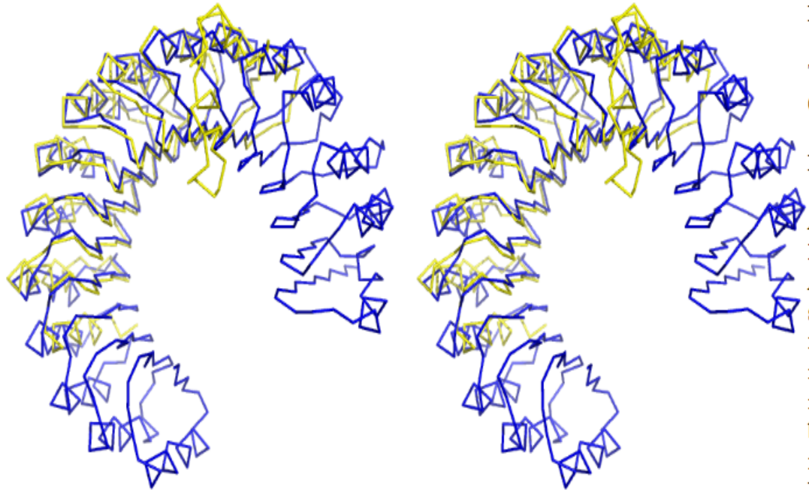

BLUE COLOR - DALI TEMPLATE

YELLOW COLOR-QUERY DOMAIN

DALI TEMPLATE ID - **1z7x-W**
DOMAIN LENGTH- **226-477 (251)**
DALI TEMPLATE LENGTH- **460**

The number of structurally equivalent residues (LALI) - **227 aa**

RMSD (DALI) - **2.3**

ANNOTATION - **LRR - Ribonuclease Inhibitor**
A leucine-rich repeat (LRR) is a protein structural motif that forms an α/β horseshoe fold. Leucine-rich repeats are frequently involved in the formation of protein–protein interactions.Leucine-rich repeat motifs have been identified in a large number of functionally unrelated proteins
Ribonuclease inhibitors (RI) or Ribonuclease/angiogenin inhibitor are a family of large proteins that bind to and inhibit ribonucleases.

*Supplementary figure 3a : Conserved domain structure of IndInt g22432*

**g22212.t1**

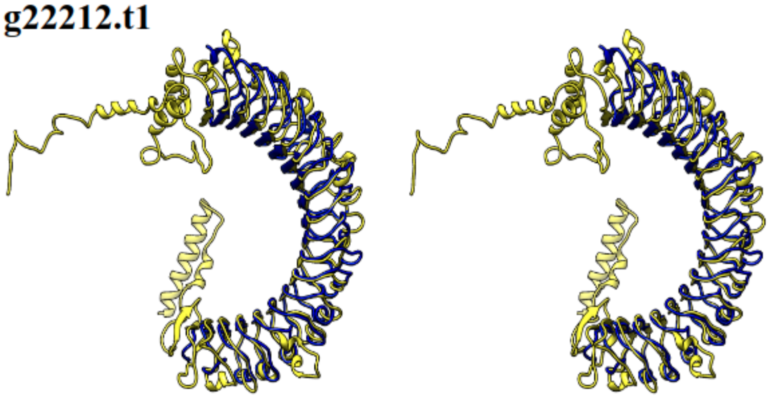

DOMAIN 1 389-1021
(635 amino acid)

BLUE COLOR - DALI TEMPLATE

YELLOW COLOR-QUERY DOMAIN

DALI TEMPLATE ID - **4li1-B**

DOMAIN LENGTH- **635 AA**

LALI - **418 AA**

DALI TEMPLATE LENGTH- **425**

RMSD (DALI) - **2.6**

Function :

LEUCINE-RICH REPEAT-
CONTAINING G-PROTEIN COUPLED

# g22212.t1

DOMAIN 2 → 435-932
(497 amino acid)

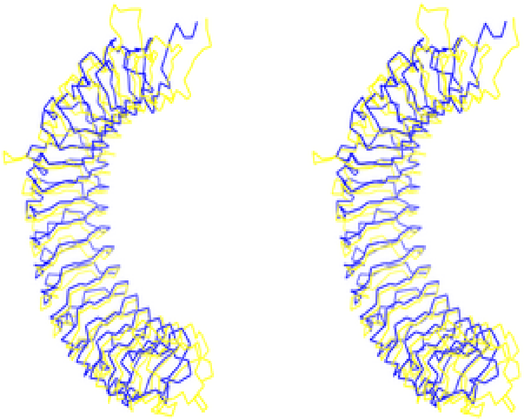

BLUE COLOR - DALI TEMPLATE

YELLOW COLOR-QUERY DOMAIN

DALI TEMPLATE ID - **4li1-B**

DOMAIN LENGTH- **497 AA**

LALI - **414 AA**

DALI TEMPLATE LENGTH- **425**

RMSD (DALI) - **2.5**

Function :

LEUCINE-RICH REPEAT-CONTAINING G-PROTEIN COUPLED

*Supplementary figure 3b : Conserved domain structure of IndInt g22212*

**g22118.t1**

BLUE COLOR - DALI TEMPLATE

YELLOW COLOR-QUERY DOMAIN

DALI TEMPLATE ID - **1JB4-B**
DOMAIN LENGTH- 181-347 (**167 AA)**
DALI TEMPLATE LENGTH-**123 AA**

The number of structurally equivalent residues (LALI) - **120 AA**

RMSD (DALI) - **2.2**

ANNOTATION - **NTF2**
[**NTF2** is a cytosolic protein responsible for nuclear import of Ran, a small Ras-like GTPase involved in a number of critical cellular processes, including cell cycle regulation, chromatin organization during mitosis, reformation of the nuclear envelope following mitosis, and controlling the directionality of nucleocytoplasmic transport]

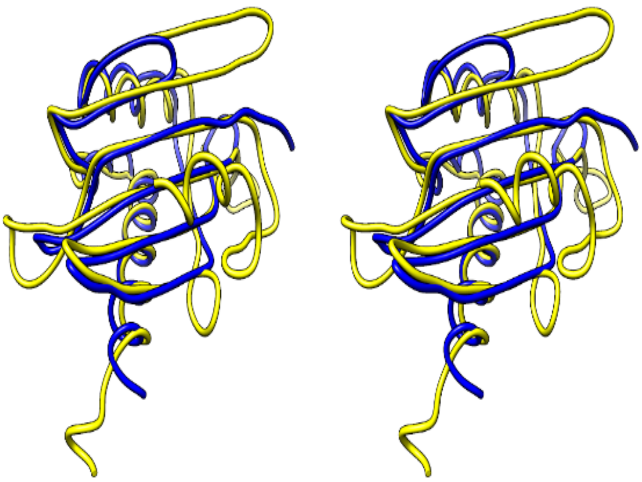

*Supplementary figure 3c : Conserved domain structure of IndInt g22118*

**g22089.t1**

BLUE COLOR - DALI TEMPLATE

YELLOW COLOR-QUERY DOMAIN

DALI TEMPLATE ID - **5MBX-A**
DOMAIN LENGTH- 1-508 **(508 AA)**
DALI TEMPLATE LENGTH- **472 AA**

The number of structurally equivalent residues(LALI)- **433 AA**

RMSD (DALI) - **2.4**

ANNOTATION-
**PEROXISOMAL N1-(ACETYL)-SPERMINE/SPERMIDINE OXIDASE**

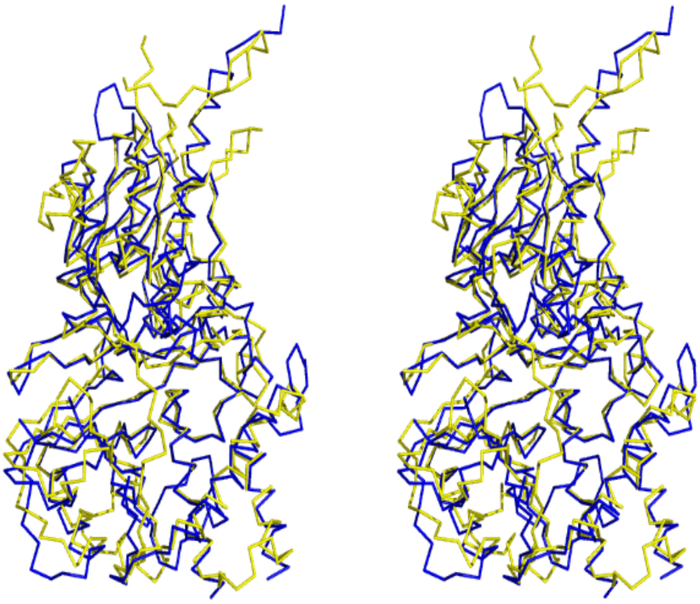

*Supplementary figure 3d : Conserved domain structure of IndInt g22089*

**g22220.t1**
**(Domain 3)**

BLUE COLOR - DALI TEMPLATE

YELLOW COLOR-QUERY DOMAIN

DALI TEMPLATE ID - **4J19-B**
DOMAIN LENGTH- 975-1042(**68 AA)**
DALI TEMPLATE LENGTH- **76 AA**

The number of structurally equivalent residues(LALI) - **65 AA**
RMSD (DALI) - **2.5**

ANNOTATION - **HOMEOBOX-CONTAINING PROTEIN 1**
[Binding to double-stranded telomere-associated DNA]

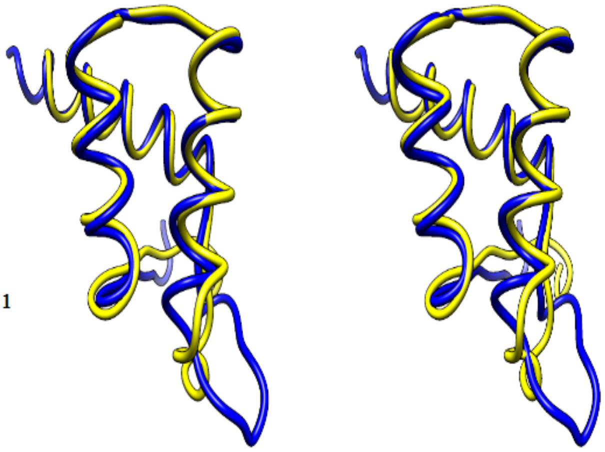

*Supplementary figure 3e : Conserved domain structure of IndInt g22220*

**g22349.t1**

DOMAIN →153-405
(252 AMINO ACIDS)

BLUE COLOUR →DALI TEMPLATE

YELLOW COLOUR → QUERY DOMAIN

DALI TEMPLATE ID - **3I0P**

DOMAIN LENGTH- **252 AA**

LALI - **253 AA**

DALI TEMPLATE LENGTH- **361 AA**

RMSD (DALI) - **1.7**

FUNCTION - MALATE DEHYDROGENASE

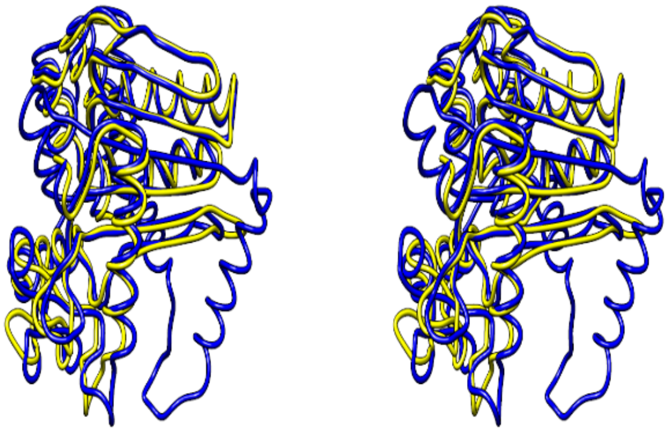

*Supplementary figure 3f : Conserved domain structure of IndInt g22349*

## g23051.t1

DOMAIN →189-468
(279 AMINO ACIDS)

BLUE COLOUR →DALI TEMPLATE

YELLOW COLOUR → QUERY DOMAIN

DALI TEMPLATE ID - **3SZ4**

DOMAIN LENGTH- **279 AA**

LALI - **180 AA**

DALI TEMPLATE LENGTH- **195 AA**

RMSD (DALI) - **2.6**

FUNCTION - EXONUCLEASE

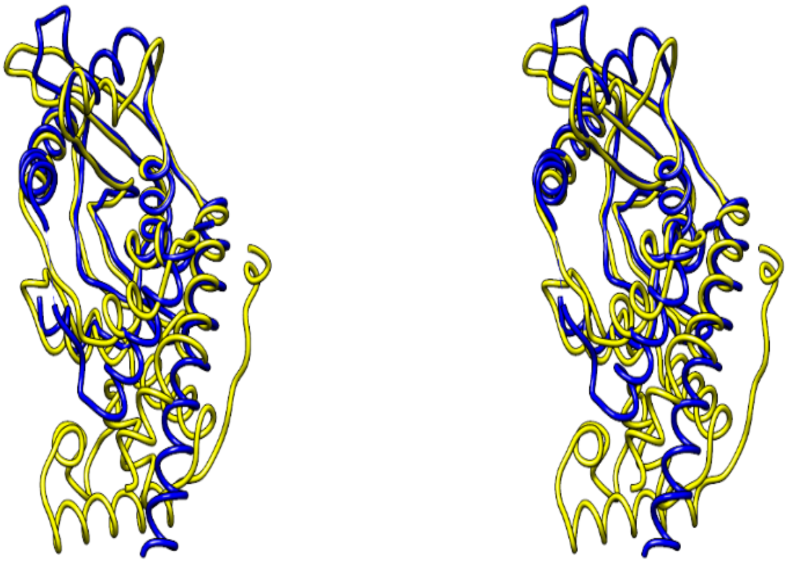

*Supplementary figure 3g : Conserved domain structure of IndInt g23051*

## g22565.t1

BLUE COLOR - DALI TEMPLATE

YELLOW COLOR-QUERY DOMAIN
DALI TEMPLATE NAME - **TUMOR NECROSIS FACTOR-INDUCIBLE GENE 6 PROTEIN**

DALI TEMPLATE ID - **2wno-A**
DOMAIN LENGTH - **128 aa**
DALI TEMPLATE LENGTH- **119 aa**

LALI - **110**

RMSD (DALI) - **2.0**

Percentage Coverage - **92.4%**
Sequence identity - **20%**

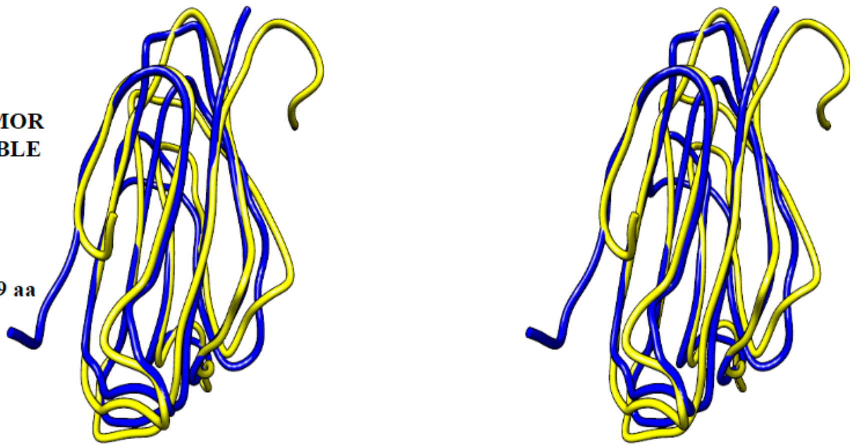

## g22565.t1

BLUE COLOR - DALI TEMPLATE

YELLOW COLOR-QUERY DOMAIN
DALI TEMPLATE NAME - **COMPLEMENT C1R SUBCOMPONENT**

DALI TEMPLATE ID - 6f1d-A
DOMAIN LENGTH - 158
DALI TEMPLATE LENGTH- 117 aa

LALI - 112

RMSD (DALI) - **2.5**

Percentage Coverage ~ 71%
Sequence identity - 24%

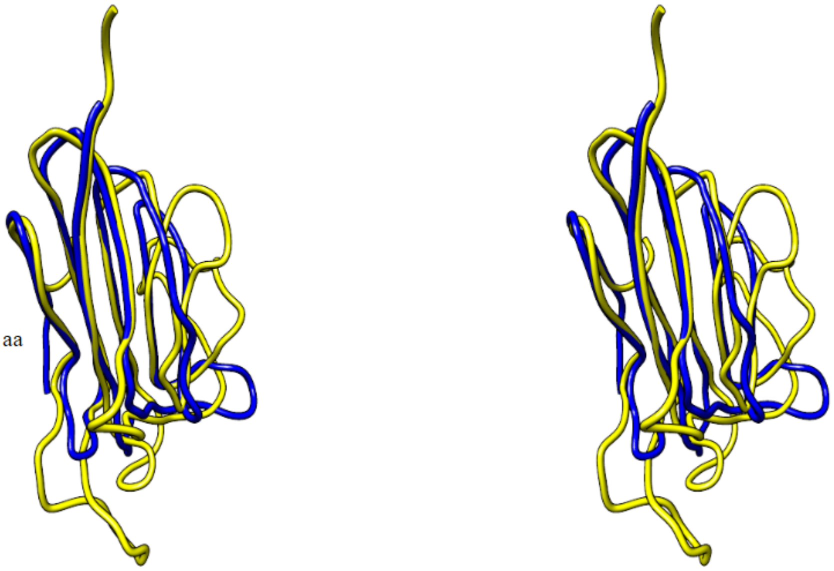

*Supplementary figure 3h : Conserved domain structure of IndInt g22565*

## g21982.t1

BLUE COLOR - DALI TEMPLATE

YELLOW COLOR-QUERY DOMAIN
DALI TEMPLATE NAME - **UBIQUITIN**
DALI TEMPLATE ID - **2j7q-A**
DOMAIN LENGTH - 248
DALI TEMPLATE LENGTH- **232**
LALI - **177**
RMSD (DALI) - **3.1**

Percentage Coverage ~ 71.4%
Sequence identity - 24%

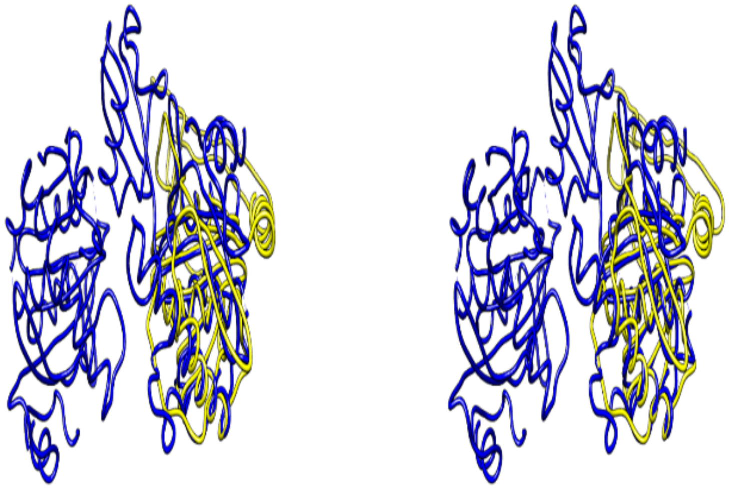

*Supplementary figure 3i : Conserved domain structure of IndInt g21982*

**g22147.t1**

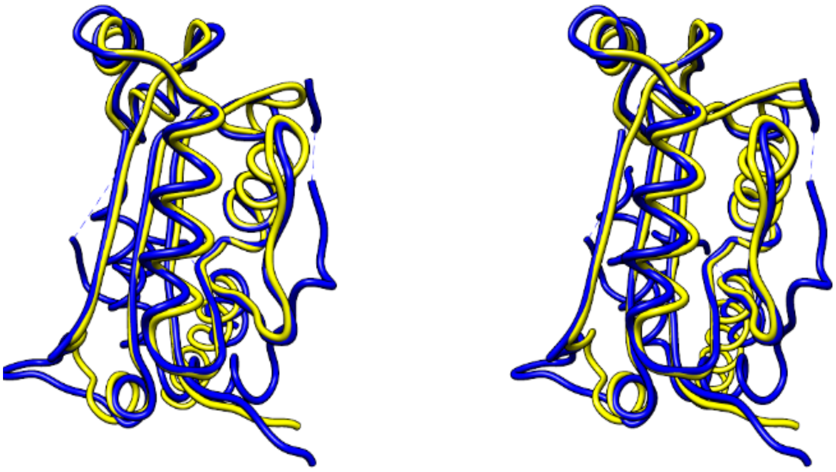

BLUE COLOR - DALI TEMPLATE

YELLOW COLOR-QUERY DOMAIN

DALI TEMPLATE ID - **1y97-B**
DOMAIN LENGTH- **1-156 (156)**
DALI TEMPLATE LENGTH- **211**

The number of structurally equivalent residues (LALI) - **145 aa**

RMSD (DALI) - **1.4**

ANNOTATION - **TREX2**
Exonuclease with a preference for double-stranded DNA with mismatched 3' termini. May play a role in DNA repair. Exonucleolytic cleavage in the 3'- to 5'-direction to yield nucleoside 5'-phosphates.

*Supplementary figure 3j : Conserved domain structure of IndInt g22147*

**g22088.t1**

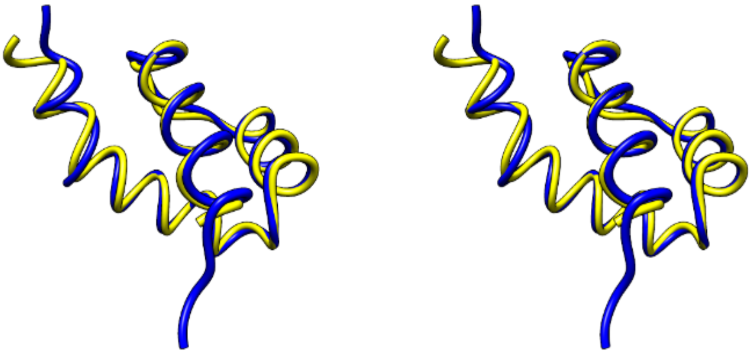

BLUE COLOR - DALI TEMPLATE

YELLOW COLOR-QUERY DOMAIN

DALI TEMPLATE ID - **6FQQ-B**
DOMAIN LENGTH- **240-297 (57)**
DALI TEMPLATE LENGTH- **61**

The number of structurally equivalent residues (LALI) - **57 aa**

RMSD (DALI) - **1.4**

ANNOTATION-**HOMEOBOX PROTEIN TGIF1**
A member of the three-amino acid loop extension (TALE) superclass of atypical homeodomains. TALE homeobox proteins are highly conserved transcription regulators. Both TGIF1 and TGIF2 act as transcription factors repressing TGF-β signalling.

*Supplementary figure 3k : Conserved domain structure of IndInt g22088*

**g22941.t1**

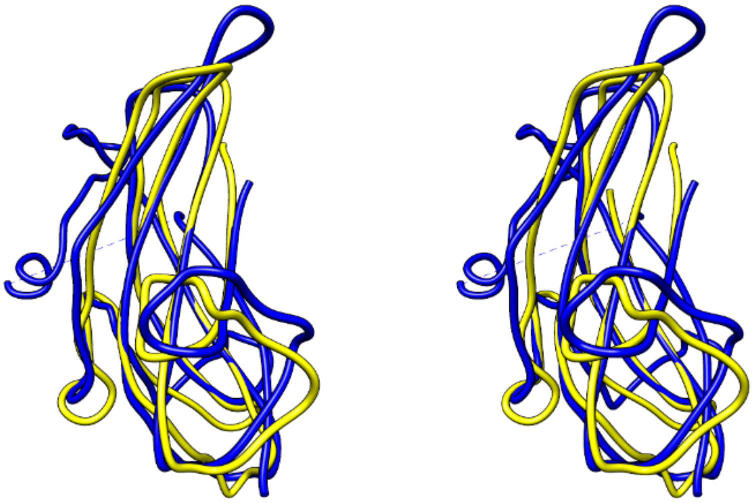

BLUE COLOR - DALI TEMPLATE

YELLOW COLOR-QUERY DOMAIN

DALI TEMPLATE ID - **4ajv-A**
DOMAIN LENGTH- **404-517 (113)**
DALI TEMPLATE LENGTH- **152**

The number of structurally equivalent residues (LALI) - **106 aa**

RMSD (DALI) - **2.1**

ANNOTATION - **TRANSFORMING GROWTH FACTOR BETA RECEPTOR TYPE 3**
Transforming growth factor beta (TGFβ) receptors are single pass serine/threonine kinase receptors that belong to TGFβ receptor family. TGFβ is a growth factor and cytokine involved in paracrine signalling and can be found in many different tissue types.

*Supplementary figure 3l : Conserved domain structure of IndInt g22941*

**g22536.t1**

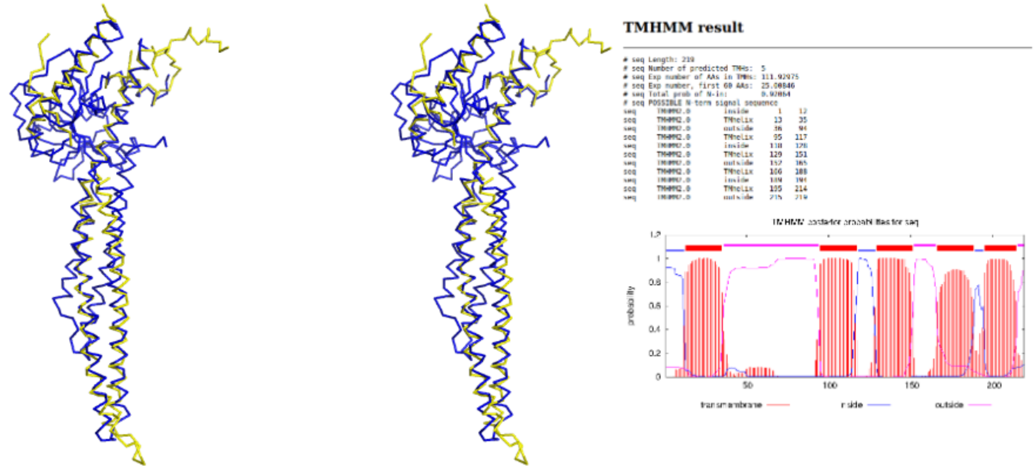


BLUE COLOR - DALI TEMPLATE

YELLOW COLOR-QUERY DOMAIN

DALI TEMPLATE ID - **6jfm-A**
DOMAIN LENGTH- **1-219 (219)**
DALI TEMPLATE LENGTH- **422**

The number of structurally equivalent residues (LALI) - **121**

RMSD (DALI) - **3.1**

ANNOTATION - **Mitofusin**
MFN2, an outer mitochondrial membrane GTPase, is critical for mitochondrial fusion, which in turn affects mitochondrial dynamics, distribution, quality control, and function.MFN2 modulates ER–mitochondria tethering.

*Supplementary figure 3m : Conserved domain structure of IndInt g22536*

**G22843**

BLUE COLOR - DALI TEMPLATE
YELLOW COLOR-QUERY DOMAIN

DALI TEMPLATE ID - **6FP9**
DOMAIN LENGTH- **209 AA**
DALI TEMPLATE LENGTH-**171 AA**
The number of structurally equivalent residues (LALI) - **168 AA**
RMSD (Chimera) - **1.897**
ANNOTATION - DARPin

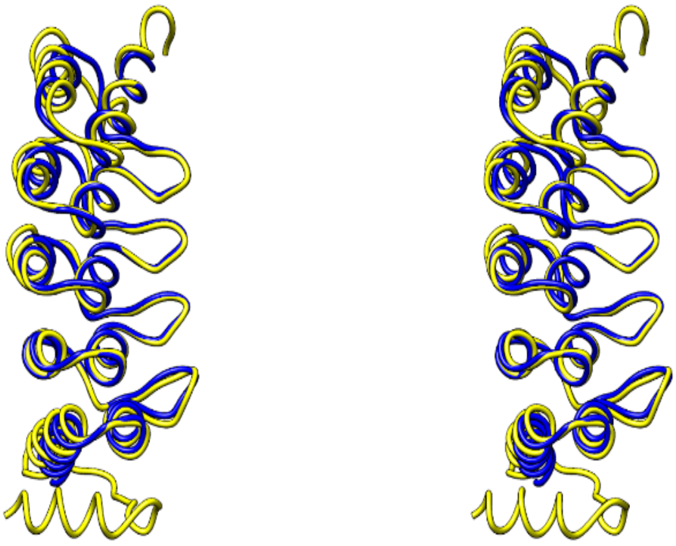

*Supplementary figure 3n : Conserved domain structure of IndInt g22843*

**G22150**

BLUE COLOR - DALI TEMPLATE
YELLOW COLOR-QUERY DOMAIN

**DALI TEMPLATE ID - 5WU1**
**DOMAIN LENGTH- 727 AA**
**DALI TEMPLATE LENGTH-493 AA**
The number of structurally equivalent residues (**LALI**) - **353 AA**
**RMSD** (Chimera) - **2.5**

**ANNOTATION -**

SPECKLE TARGETED PIP5K1A-REGULATED POLY(A) POLYMERASE.
Poly(A) polymerase that creates the 3'-poly(A) tail of specific pre-mRNAs

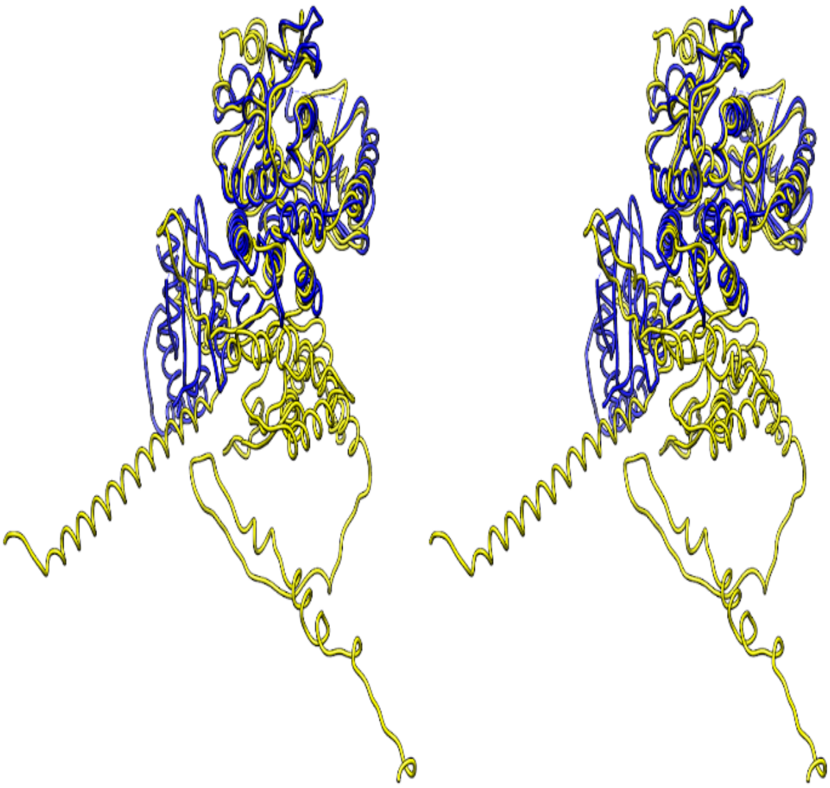

*Supplementary figure 3o : Conserved domain structure of IndInt g22150*

**DOMAINS WHICH DID NOT SUPERIMPOSE WELL BUT HAVE A FUNCTIONAL ANNOTATION IN DALI**

GENE ID - **g22118.t1**
DOMAIN LENGTH - 1-109(**109 AA**)

FIRST HIT DALI TEMPLATE PDB ID - **7K3R-A**
DALI TEMPLATE LENGTH- **95 AA**

The number of structurally equivalent residues(LALI) - **80 AA**

RMSD (DALI) - **3.5**

ANNOTATION - **INTERFERON INDUCIBLE PROTEIN AIM2**

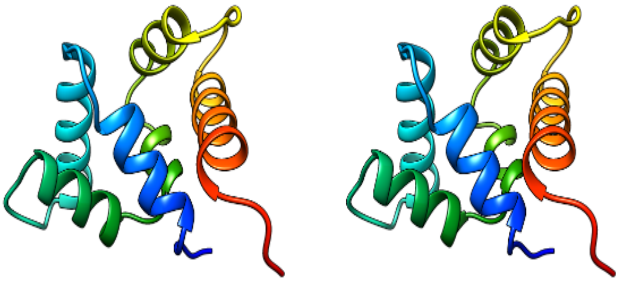

*Supplementary figure 3p : DALI functional annotation of IndInt g22118*

**G22810.t1 (not superimposed well with template)**

DOMAIN 1
246-561
(315 amino acid)

DALI TEMPLATE ID - **7lw7-A (only one hit )**

DOMAIN LENGTH- **315 AA**

LALI - **80 AA**

DALI TEMPLATE LENGTH- **277**

RMSD (DALI) - **5.7**

Function :
EXONUCLEASE V
The Exonuclease V enzyme is an ATP-dependent, double-strand DNA exonuclease.

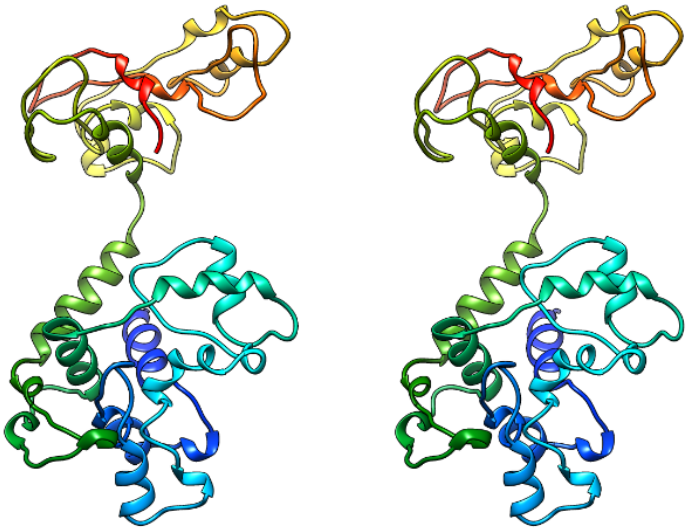

*Supplementary figure 3q : DALI functional annotation of IndInt g22810*

**g22301.t1 (not superimposed well with template)**

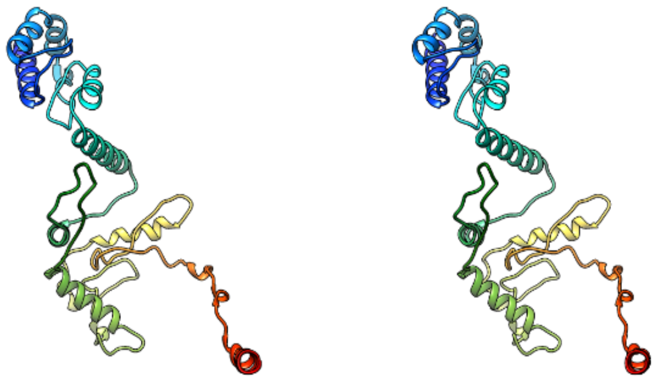

DOMAIN 1
21-306
(285 amino acid)

DALI TEMPLATE ID - **2v71-A**
DOMAIN LENGTH- **285 AA**

LALI - **156 AA**

DALI TEMPLATE LENGTH- **160**

RMSD (DALI) - **5.2**

Function :
NUCLEAR DISTRIBUTION PROTEIN NUDE-LIKE 1
It plays a role in multiple processes including cytoskeletal organization, cell signaling and neuron migration, outgrowth and maintenance

*Supplementary figure 3r : DALI functional annotation of IndInt g22301*

**g23001 (not superimposed properly)**

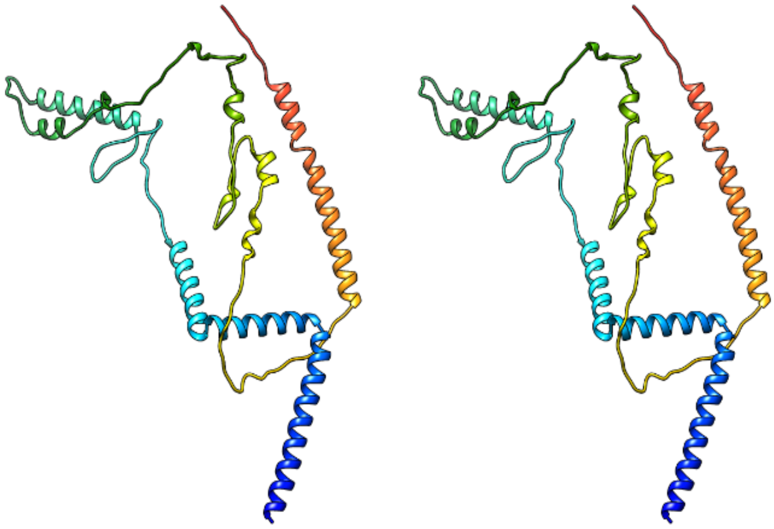

DOMAIN
115-395
(280 amino acid)

DALI TEMPLATE ID - **1xpj-D**
DOMAIN LENGTH- **280 AA**

LALI - **106 AA**

DALI TEMPLATE LENGTH- **123**

RMSD (DALI) - 2.6

The Hit template is a HYPOTHETICAL PROTEIN;

*Supplementary figure 3s : DALI functional annotation of IndInt g23001*

### MODELS WHICH DID NOT FOLD WELL-

**MODELS WHICH DID NOT FOLD WELL**

**g22846.t1(length - 701 AA)**

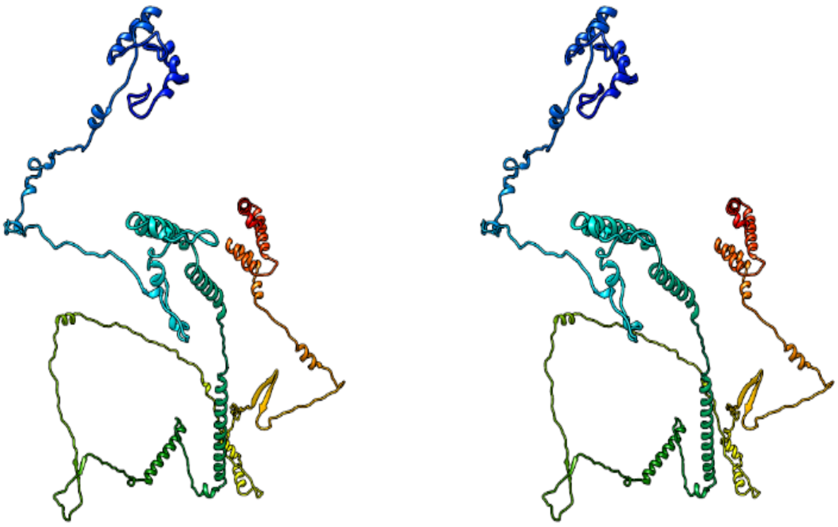

*Supplementary figure 3t : Alpha fold predicted structure of IndInt g22846*

**g22224.t1 (LENGTH - 343 AA)**

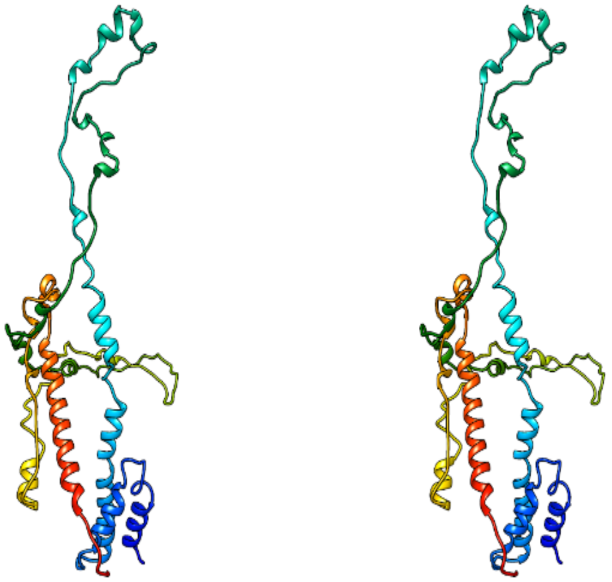

*Supplementary figure 3u : Alpha fold predicted structure of IndInt g22224*

**g22323 450 amino acids - very high error plot**

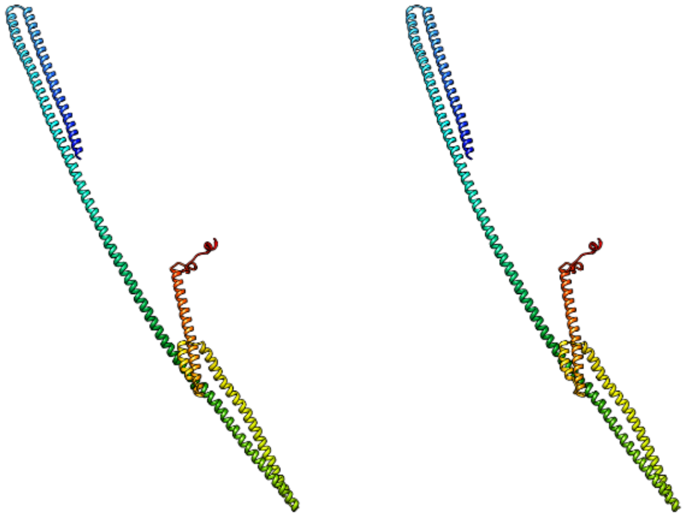

*Supplementary figure 3v : Alpha fold predicted structure of IndInt g22323*

**g21978.t1**

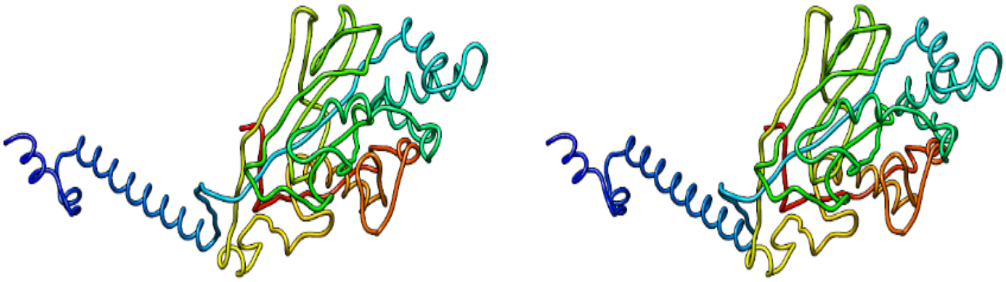

*Supplementary figure 3w : Alpha fold predicted structure of IndInt g21978*

**g22197.t1**

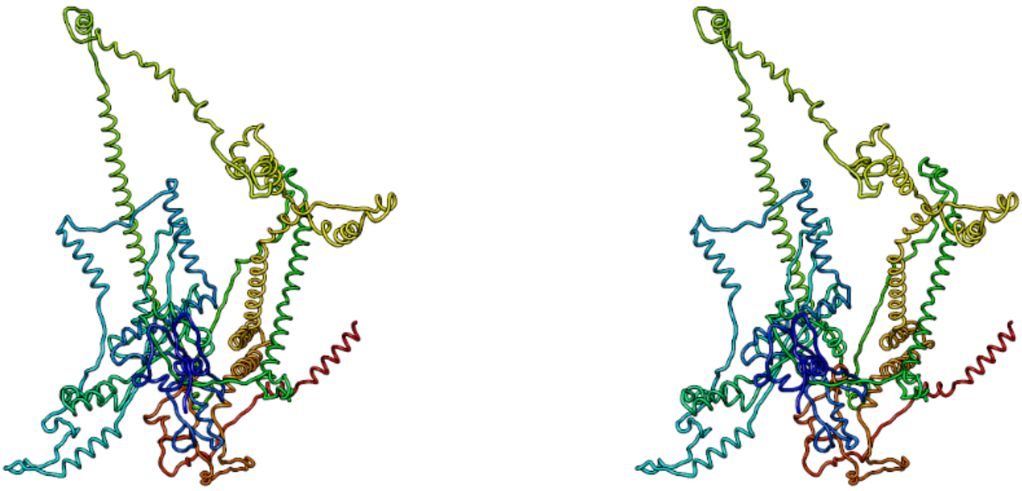

*Supplementary figure 3x : Alpha fold predicted structure of IndInt g22197*

**g22416.t1**

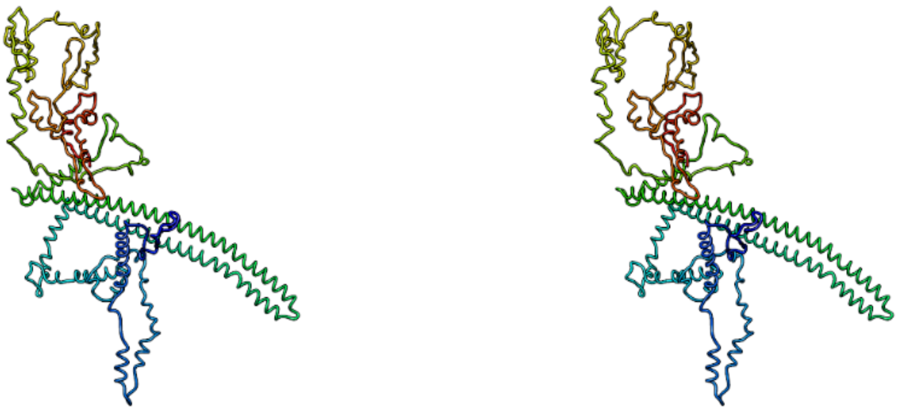

*Supplementary figure 3y : Alpha fold predicted structure of IndInt g22416*

**g22689.t1**

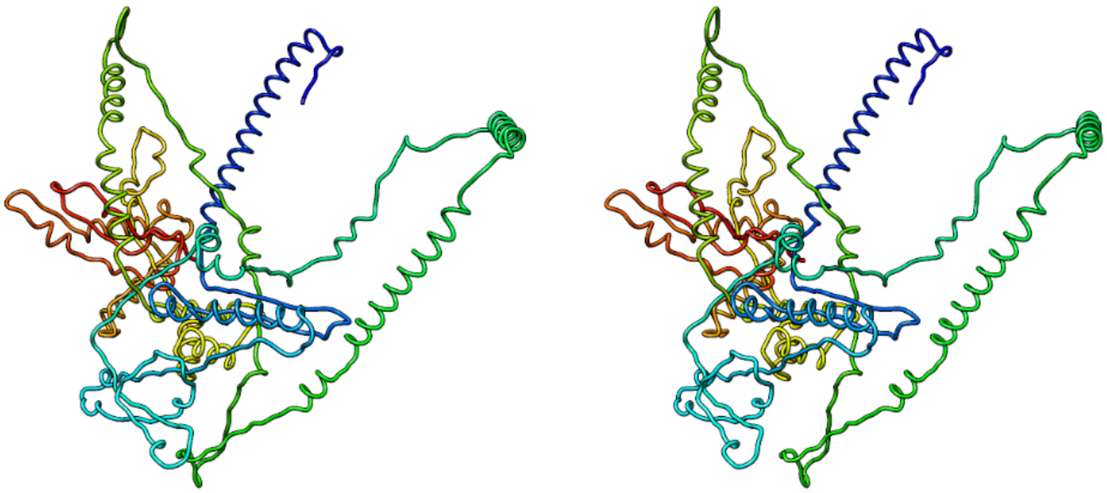

*Supplementary figure 3z : Alpha fold predicted structure of IndInt g22689*

**g21989.t1**

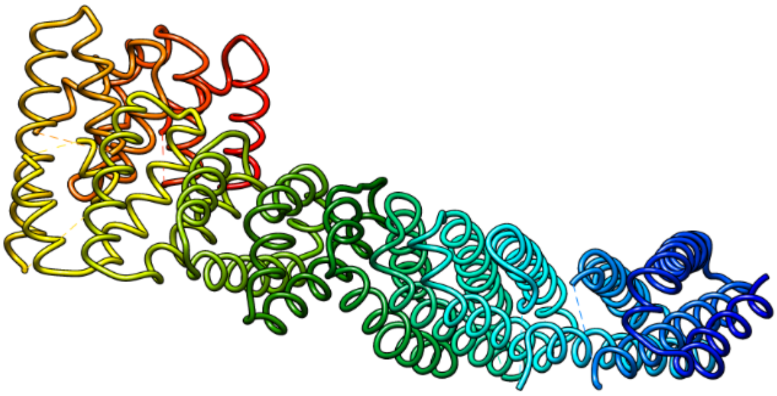

*Supplementary figure 3a1 : Alpha fold predicted structure of IndInt g21989*

**g22713.t1**

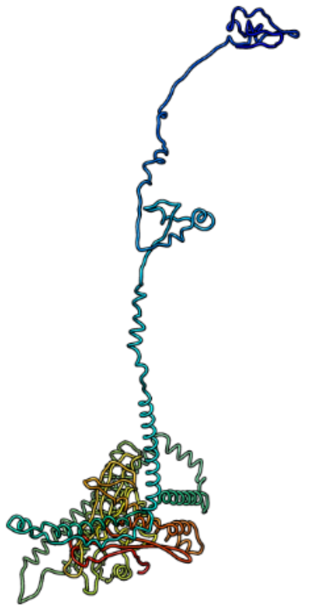

*Supplementary figure 3a2 : Alpha fold predicted structure of IndInt g22713*

**g23157.t1**

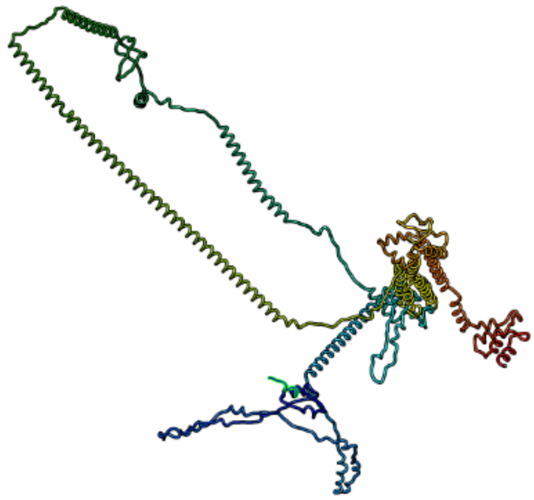

*Supplementary figure 3a3 : Alpha fold predicted structure of IndInt g23157*

**g22302.t1**

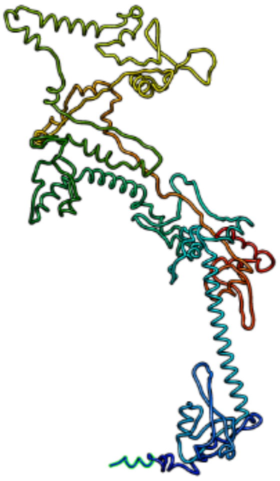

*Supplementary figure 3a4 : Alpha fold predicted structure of IndInt g22302*

**g22313.t1**

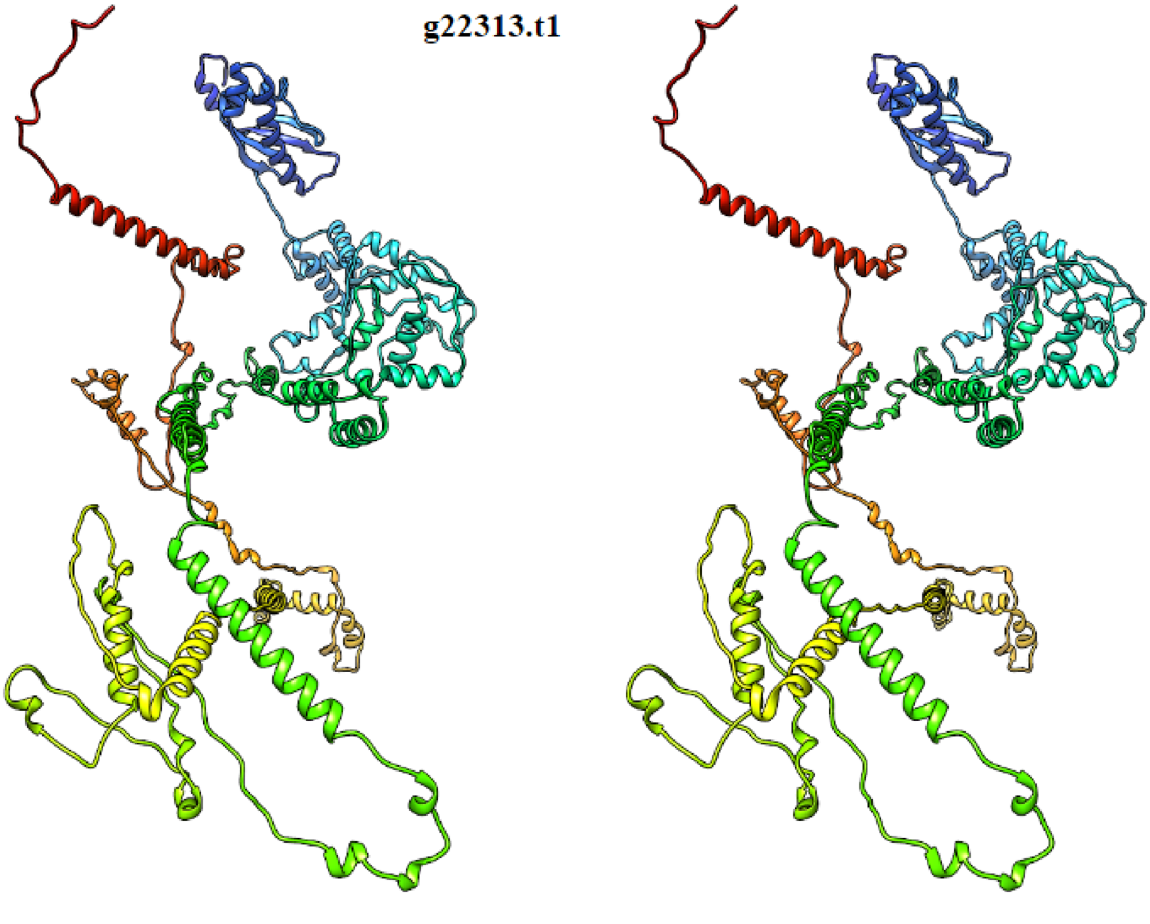

*Supplementary figure 3a5 : Alpha fold predicted structure of IndInt g22313*

```
CLUSTAL O(1.2.4) multiple sequence alignment


AnColuzzii_XP040229266        -----------------------------------------------MSDTNLLQLSAAW    13
AnArabiensis_XP040157989      -----------------------------------------------MSDTNLLQLSAAW    13
AnGambiae_AGAP005805          -----------------------------------------------MSDTNLLQLSAAW    13
IndInt_g22432                 -----------------------------------------------MSDTNLLHLSALW    13
AnStephensi_XP035917226       MFRISHATDSFDPAADVRIIEALKLLHQVQTEPSKGIAQPISCERIKMSDTNLLHLSALW    60
                                                                             *******:*** *

AnColuzzii_XP040229266        TQLPRLMVPFDRQLSNALRTWAPSWQTQELASETTRKLFVELVTFGGVNQCRRFDALARP    73
AnArabiensis_XP040157989      RQLPRLMVPFDRQLSNALRTWAPSWQTQQLASETTRKLFVELVTFGGVNQCRRFDALARP    73
AnGambiae_AGAP005805          RQLPRLMVPFDRQLSNALRTWAPSWQTQQLASETTRKLFVELVTFGGVNQCRRFDALARP    73
IndInt_g22432                 RQLPHLTSPLERELWNAFREYGPSWQTQQLASETQRELFGELVAFGGRRACRPFDAMVHV    73
AnStephensi_XP035917226       RQLPHLTSPLERELWNAFRAYGPSWQTQQLASETQRELFVELVAFGGRRTCRPFDAMVHV    120
                               ***:*  *::*:* **:* :.******:***** *:** ***:*** . ** ***:.:

AnColuzzii_XP040229266        VINDHGEGVVPPRLSTIVVQFLVTRYDGGPEPTALSCCQVRDFGCLLSTELPLLKIVDLE    133
AnArabiensis_XP040157989      VINDHGEGVVPARLSTIVVQFLVARYDGGPEPAALSCCQVREFGCLLSTELPLLKIVDLE    133
AnGambiae_AGAP005805          VINDHGEGVVPPRLSTIVVQFLVTRYDGGPEPTALSCCQVREFGCLLSTELPLLKIVDLE    133
IndInt_g22432                 --TDDIVQDVPKRLSTLVLEFLVASFDGGPEPGPLNCCQVREFGSLLSTELPLIKVIELE    131
AnStephensi_XP035917226       --TDDIVQDVPKRLSTLVLEFLVVSFDGGPEPGPLSCCQVREFGSLLSTELPLIKVIELE    178
                                .*.    ** ****:*::***. :******  *.*****:**.********:*:::**

AnColuzzii_XP040229266        SETYWRRVVWCYTDEPLSYYEHDYLTPHRFWKQHGVELKLARMIEQQDPMYWELEGLEDT    193
AnArabiensis_XP040157989      SETYWRRVVWCYADEPLSYYEHDYLTPRRFWKQHGIELKLARMIEQQDPVYWELEGLEDT    193
AnGambiae_AGAP005805          SETYWRRVVWCYADEPLSYYEHDHLTPRRFWKQHGVELKLARMIEQQDPVYWELEGLEDT    193
IndInt_g22432                 SETYWRRVVWCYSNDSLSYHELDYLSPRRFWKQQGIELKLARMIEQQDPQYWELEGLEET    191
AnStephensi_XP035917226       SETYWRRVVWCYSNDSLSYHELDYLSPRRFWKQQGIELKLARMIEQQDPHYWELEGLEET    238
                              ************::: ***:* *:*:*:*****:*:************* ********:*

AnColuzzii_XP040229266        IKRAASFVNNLVVQQLRPYAMVEPLEDYEAYTIRNAPADLCHHGSLAVLGHLVNLTSLSL    253
AnArabiensis_XP040157989      IKRAAPFVNNLVVQQLRPYAMVEPLEDYEAYTIRNAPADLCHHGSLAVLGHLVNLTSLSL    253
AnGambiae_AGAP005805          IKRAAPFVNNLVVQQLRPYAMVEPLEDYEAYTIRNAPADLCQHGSLAVLGHLVNLTSLSL    253
IndInt_g22432                 IKKAAPFVDTLYVEQLQPLPTVEPIEDYEEYHIRNAPDELCNHGSLAILGHLVNLTSLSL    251
AnStephensi_XP035917226       IKRAAPFVDTLYIEQLQPHPTVEPIEDYEEYHIRNAPDELCNHGSLAILGHLVNLTSLSL    298
                              **:** **:.* ::**:*   ***:**** * ***** :**:*****:************

AnColuzzii_XP040229266        VFGVKHWTKAYQTRYSQCSQTDIEQLGMGLQKLVKLKKFNISHTRLNASKLKALLESLTP    313
AnArabiensis_XP040157989      MFGVKHWTKAYQTRYSHCSQTDIEQLGMGLQKLVKLKKFNLSHTRLNASKLKPLLESLTP    313
AnGambiae_AGAP005805          VFGVKHWNKAYQTRYSHCSQTDIEQLGMGLQKLVKLKKFNLSHTRLNASKLKALLESLTP    313
IndInt_g22432                 VFGVKHWTKPYQNRFSSCSLADIENLGQGLQKLHRLKKFTLSHTQLNSAKLKILLEFLTP    311
AnStephensi_XP035917226       VFGVKHWIKPYQNRFSSCSLADIENLGQGLQKLHRLKKFTLSHTQLNSAKLKILLEFLTP    358
                              :****** * **.*:* ** :***:** ***** :****.:***:**::*** *** ***

AnColuzzii_XP040229266        LGLEAIRLTHCHLGEGCGGPLGRFLSRFGPTLKQLDLSNNRLDAPELDQLCPGLSIYQGT    373
AnArabiensis_XP040157989      LGLETIRLTHCHLGEGCGGPLGRFLSRFGPTLKQLDLSNNRLDAPELDQLCPGLSIYQGT    373
AnGambiae_AGAP005805          LGLETIRLTHCHLGEGCGGPLGRFLSRFGPTLKQLDLSNNRLDAPELDQLCPGLSIYQGK    373
IndInt_g22432                 LELETIRLTHCQLAAGCGAILGRFLSRFQPTLTQLDLSHNRLDAPELDQLCPGLSVYRGV    371
AnStephensi_XP035917226       LELETIRLTHCQLAAGCGAILGRFLSRFQPTLTHLDLSHNRLDAPELDQLCPGLSVYRGV    418
                              * **:******:*. ***. ******** ***.:****:****************:*:*
```

```
AnColuzzii_XP040229266        LDKLNLSYNPIGEAGVLILGGAIKGRAQLSELHFTGCPMGVEGSFRVIQLLSFHETLRKV    433
AnArabiensis_XP040157989      LDKLNLSYNPIGEAGVLILGGAIKGRAQLSELHFTGCPMGVEGSFRVIQLLSFHETLRKV    433
AnGambiae_AGAP005805          LDKLNLSYNPIGQAGVLILGGAIKGRAQLSELNFTGCPLGVEGSFRVIQLLSFHETLRKV    433
IndInt_g22432                 VDRLDLSYNPIGEAGVLILGGAIKGRTQLSELNFTGCQMGVEGSFRL---LSFHVTLRKV    428
AnStephensi_XP035917226       VDRLDLSYNPIGEAGVLILGGAIKGRTQLSELNFTGCQMGVEGSFRVIQLLSFHVTLRKI    478
                              :*:*:*******:*************:*****:**** :*******:   **** ****:

AnColuzzii_XP040229266        SLNCVPISPEGGDKLVQVLLENNRIEEVQVRDCGLTERVRSKIGKILRKNASTRHQSRRS    493
AnArabiensis_XP040157989      SLNCVPISPEGGDKLVQVLLENNRIEEVQVRDCGLTERVRARIGKILRKNANARHRSRRS    493
AnGambiae_AGAP005805          SLNCVPISPEGGDKLVQVLLENNRIEEVHVRDCGLTERVRVRIGKILRKNANARHRSRRS    493
IndInt_g22432                 SLNCVPISLEGGAKLVQVLQENIRIEDVQVRQCGLPDELLVKPHIIFANPDYRASSSGDT    488
AnStephensi_XP035917226       SLNCVPISLEGGAKLVQVLQENIRIEDVQVRQCGLPDELLVKVRKILRKNAKIRDQLR-S    537
                              ******** *** ****** ** ***:*:**:*** :.:  :   *: :          :

AnColuzzii_XP040229266        DGKVATPPMVRSPAT-------------------MFQLERQVFPVQS-------------    521
AnArabiensis_XP040157989      DGKVATPPMVRSPAS-------------------MFQLERQVFPVQS-------------    521
AnGambiae_AGAP005805          AGKVATPPMVRSPAS-------------------MFQLERQVFPVQS-------------    521
IndInt_g22432                 ALLTAPKEIRHCPPVDRQAEPVPGRTIIHYANKSHFPFPVRILVKVLRPVAFSGPTRTKL    548
AnStephensi_XP035917226       VRMSSEQPMDKCHTSERSLGS-------------MFLLERQ----------VFSE------    569
                                  :   : :.                       * :  :

AnColuzzii_XP040229266        ------------------------------------------------------------    521
AnArabiensis_XP040157989      ------------------------------------------------------------    521
AnGambiae_AGAP005805          ------------------------------------------------------------    521
IndInt_g22432                 CRVAPGPTFASASDSEQTLALWSECNTTRADRSTADRPIRSRSIGGLIVKWECLWHGPEH    608
AnStephensi_XP035917226       ------------------------------------------------------------    569


AnColuzzii_XP040229266        ------------------------------------------------------------    521
AnArabiensis_XP040157989      ------------------------------------------------------------    521
AnGambiae_AGAP005805          ------------------------------------------------------------    521
IndInt_g22432                 KRMCACASVCVLCMWAPRFGRHFDGSMTLKTSTGNIYQTGYRLSGTSGRITSACFTRVHP    668
AnStephensi_XP035917226       ------------------------------------------------------------    569


AnColuzzii_XP040229266        ------------------------- 521
AnArabiensis_XP040157989      ------------------------- 521
AnGambiae_AGAP005805          ------------------------- 521
IndInt_g22432                 GTICSGFMATGERHTHAHIPPTTRK 693
AnStephensi_XP035917226       ------------------------- 569
```

*Supplementary figure 4a : MSA for IndInt gene g22432.t1 across other Anopheles species shows high conserved homology.*

```
CLUSTAL O(1.2.4) multiple sequence alignment


AnColuzzii_XP040230463          ------------------------------------------------------------      0
AnArabiensis_XP040158968        ------------------------------------------------------------      0
AnGambiae_AGAP006645            ------------------------------------------------------------      0
AnStephensi_XP035917013         ------------------------------------------------------------      0
IndInt_g22212                   MRGLTRSVDPTARLQDHETERTVVCAYGYIGIPDISFWQLVNAPLRESEMASEPYTVPDS      60

AnColuzzii_XP040230463          --------------------------------------------------MVSVHTMRCQ      10
AnArabiensis_XP040158968        --------------------------------------------------MVSVHTMRCQ      10
AnGambiae_AGAP006645            --------------------------------------------------------MRCQ      4
AnStephensi_XP035917013         --------------------------------------------------------MSRQ      4
IndInt_g22212                   GCMAMHSRKSVEECRVQSRNDGGFSPARMQTPPWYNICSNLTVPLIFFTQQGSIHTMSRQ      120
                                                                                        *  *

AnColuzzii_XP040230463          YLLVLTVVLCALTVN---SLKPDLPKAKDPLFNKMLAESSNKGVDAQDPRYNKMLPDLEE      67
AnArabiensis_XP040158968        YLLVLTVVLCALTVN---SLKPDLPKAKDPLFNKMLAESSTKGVDVQDPRYNKMLPDLEE      67
AnGambiae_AGAP006645            YLLVLTVVLCALTVN---SLKPDLPKAKDPLFNKMLAESSTKGVDAQDPRYNKMLPDLEE      61
AnStephensi_XP035917013         YLLVLAVALCALAVHSSPALKPDLPKAQDPLYNKILAETTRSVMDAHDRHYNKMLPQPDE      64
IndInt_g22212                   YLLVLAVALCALAVNSSPALKPDLPKAQDPLYNKILAETTRSVMDAHDRHYNKMLPQPDE      180
                                *****:*.****:*:   :********:***:**:***:: . :*.:* :******: :*

AnColuzzii_XP040230463          NLIDDDDEDDDEEEEDDV----APPTKKAADLNPMYALRGKQ-VKPEVSTVGSVAINKVK      122
AnArabiensis_XP040158968        NLIDDDDEDDDEEEEDDV----APPTKKAADLNPMYALRGKQ-VKPEVSTVGSVAINKVK      122
AnGambiae_AGAP006645            NLIDDDDEDDDEEEEDDV----APPTKKAADLNPMYALRGKQ-VKPEVSTVGSVAINKVK      116
AnStephensi_XP035917013         NLINSDISEDEDGDEDDAMANDDILPKKAADLNPMYAQRGKQPAKPEVSTVGSVAINKVK      124
IndInt_g22212                   NLINSDISEDEDADDDEAMANDDILPKKAADLNPMYAQRGKQPAKPEVSTVGSVAINKVK      240
                                ***:.* .:*:: ::*:.        *********** **** .****************

AnColuzzii_XP040230463          INEDIDSYEEVLLKGNGKSV--------------KPSKSTTAKPVAKPATEDNYDEYDDD      168
AnArabiensis_XP040158968        INEDIDSYEEVLLKGNGKSV--------------KPSKSTTAKPVAKPATEDNYDEYDDD      168
AnGambiae_AGAP006645            INEDIDSYEEVLLKGNGKSVDIDDSYQVEDLSGEKPSKSTTAKPVAKPATEDNYDEYDDD      176
AnStephensi_XP035917013         INEDIDSYEEVLLKGNGKSV--------------KPSKPTTSKPAAKTEGED--DE-YDD      167
IndInt_g22212                   INEDIDSYEEVLLKGNGKSV--------------KPSKPTTSKPAAKTEGED--DE-YDD      283
                                ********************              **** **:**.**   **  **  **

AnColuzzii_XP040230463          ESDEQIDFSGVDKVLAQPLKPQVPAEKSAKPAVPSTTAKPKAVTTGKQNVEILDQYDDDA      228
AnArabiensis_XP040158968        ESDEQIDFSGVDKVLAQPLKPQVPAEKSAKPAVPSTTAKPKAVTTGKQNVEILDQYDDDA      228
AnGambiae_AGAP006645            ESDEQIDFSGVDKVLAQPLKPQVPAEKSAKPAVPSTTAKPKAVTTGKQNVEILDQYDDDA      236
AnStephensi_XP035917013         ESDEKIDFSGVDKVLAQPLKP-VPAEKSVKTSPPSTTLKPKSTTTGKQNVEILDQYDEDA      226
IndInt_g22212                   ESDEKIDFSGVDKVLAQPLKP-VPAEKSVKTSPPNTTLKPKSTTTGKQNVEILDQYDDDA      342
                                ****:**************** ******.* : *.** ***:.**************:**

AnColuzzii_XP040230463          DMDYQYNYDDDEDNDDDDDEDDDDDEEEDAETLPAQKITNKVNKEVAKKQTNPVKETKDQ      288
AnArabiensis_XP040158968        DMDYQYNYDDDEDNDDDDDEDDDDDEEEDAETLPAQKITNKVNKEVAKKQTNPVKETKDQ      288
AnGambiae_AGAP006645            DMDYQYNYDDDEDNDDDDDEDDDDDEEEDAETLPAQKITNKVNKEVAKKQTNPVKETKDQ      296
AnStephensi_XP035917013         DMDYQYTYDDDEDNDDDEDDDDD-DDDEDAEELPPQKAGKNVNSE-AKKQTKPVKKTKEQ      284
IndInt_g22212                   DMDYQYTYDDDEDNDDDEDDDDD-DDDEDAEELPPQKAGKNVNSE-AKKQTKPVKKTKEQ      400
                                ******.**********:*:*** *::**** ** **  ::**.* *****:***:**:*
```

```
AnColuzzii_XP040230463        NKAAGKDNAEDYYDDDEYDD--DDNAEPSNSSYCPRGCICERNMHAYMVATCSRLDLETQ      346
AnArabiensis_XP040158968      NKAAGKDNAEDYYDDDEYDD--DDNAEPSNSSYCPRGCICERNMHAYMVATCSRLDLETQ      346
AnGambiae_AGAP006645          NKAAGKDNAEDYYDDDEYDD--DDNAEPSNSSYCPRGCICERNMHAYMVATCSRLDLETQ      354
AnStephensi_XP035917013       NKTAAKGTDESYYYDDEDDDDDDYNVDSSNSTYCPRGCICERNMHSYMVATCSRLDLETQ      344
IndInt_g22212                 NKTAAKSTDESYYYDDEDDDEDDYNVDSSNSTYCPRGCICERNMHSYMVATCSRLDLETQ      460
                              **:*.*.. *.** *** **  * *.: ***:*************:**************

AnColuzzii_XP040230463        KFTSHITDLQVLDVGPKYPIELGPEFFKKIGLSHVVSIKITNCTIVYISPQAFAGLDELY      406
AnArabiensis_XP040158968      KFTSHITDLQVLDVGPKYPIELGPEFFKKIGLSHVVSIKITNCTIVYISPQAFAGLDELY      406
AnGambiae_AGAP006645          KFTSHITDLQVLDVGPKYPIELGPEFFKKIGLSHVVSIKITNCTIVYISPQAFAGLDELY      414
AnStephensi_XP035917013       KFTSAITDLQVLDVGPKYPIELGPEFFKKIGLSHVVSIKITNCTIVYISPQAFAGLDVLY      404
IndInt_g22212                 KFTSAITDLQVLDVGPKYPIELGPEFFKKIGLSHVVSIKITNCTIVYISPQAFAGLDVLY      520
                              **** **************************************************** **

AnColuzzii_XP040230463        SVNLTNSGIDIIHPDTFANNTKLRLLTLSGNDLSAMQSVNHNTPYMDYMLKAPTVEELDI      466
AnArabiensis_XP040158968      SVNLTNSGIDIIHPDTFANNTKLRLLTLSGNDLSAMQSVNHNTPYMDYMLKAPTVEELDI      466
AnGambiae_AGAP006645          SVNLTNSGIDIIHPDTFANNTKLRLLTLSGNDLSAMQSVNHNTPYMDYMLKAPTVEELDI      474
AnStephensi_XP035917013       SVNLTNSGIDMIHPDTFANNTKLRLLTLSGNDLSAMQSVNHNTPYMDYMLKAPTVEELDI      464
IndInt_g22212                 SVNLTNSGIDMIHPDTFANNTKLRLLTLSGNDLSAMQSVNHNTPYMDYMLKAPTVEELDI      580
                              **********:*************************************************

AnColuzzii_XP040230463        SRCKLQELQPNAFNELKNIIYINLSENNLSNLPEGIFDNVETIEELDLSMNNIVELPKNI      526
AnArabiensis_XP040158968      SRCKLQELQPNAFNELKNIIYINLSENNLSNLPEGIFDNVETIEELDLSMNNIVELPKNI      526
AnGambiae_AGAP006645          SRCKLQELQPNAFNELKNIIYINLSENNLSNLPEGIFDNVETIEELDLSMNNIVELPKNI      534
AnStephensi_XP035917013       SRCKLQELQPNAFNELKNIIYINLSENNLSNLPEGIFDNVETIEELDLSANNIAELPKNI      524
IndInt_g22212                 SRCKLQELQPNAFNELKNIIYINLSENNLSNLPEGIFDNVETIEELDLSANNIAELPKNI      640
                              ************************************************* ***.******

AnColuzzii_XP040230463        FAKTSLAILHLKHNKITNNVDFVTADLQKLDLSFCQIRTVHNTMFKGMDGLTNLILKGNH      586
AnArabiensis_XP040158968      FAKTSLAILHLKHNKITNNVDFVTADLQKLDLSFCQIRTVHNTMFKGMDGLTNLILKGNH      586
AnGambiae_AGAP006645          FAKTSLAILHLKHNKITNNVDFVTADLQKLDLSFCQIRTVHNTMFKGMDGLTNLILKGNH      594
AnStephensi_XP035917013       FAKTSLAILHLKHNKISNNVDFVTADLQKLDVSFCQIRTINNQMFKGMEGLTNLILKGNH      584
IndInt_g22212                 FAKTSLAILHLKHNKISNNVDFVTADLQKLDVSFCQIRTINNQMFKGMEGLTNLILKGNH      700
                              ****************:**************:*******::* *****:***********

AnColuzzii_XP040230463        IEKIKPMAFISLKNLRQIDLSYNNLEQISAQTFIGNKMLDIIRLNNNPRLKRLPNEGFEI      646
AnArabiensis_XP040158968      IEKIKPMAFISLKNLRQIDLSYNNLEQISAQTFIGNKMLDIIRLNNNPRLKRLPNEGFEI      646
AnGambiae_AGAP006645          IEKIKPMAFISLKNLRQIDLSYNNLEQISAQTFIGNKMLDIIRLNNNPRLKRLPNEGFEI      654
AnStephensi_XP035917013       IEKIKPMAFISLSSLRQIDLSYNNLEQISAQTFIGNKMLDIIRMNNNPRLKRLPNEGFES      644
IndInt_g22212                 IEKIKPMAFISLSSLRQIDLSYNNLEQISAQTFIGNKMLDIIRMNNNPRLKRLPNEGFES      760
                              ************..*****************************:***************

AnColuzzii_XP040230463        SFNGTFTVYFMDVSNCDISELADNTFKTMPHLTRLNLAWNNLQTIRSTYFAHLNKLMDLD      706
AnArabiensis_XP040158968      SFNGTFTVYFMDVSNCDISELADNTFKTMPHLTRLNLAWNNLQTIRSTYFAHLNKLMDLD      706
AnGambiae_AGAP006645          SFNGTFTVYFMDVSNCDISELADNTFKTMPHLTRLNLAWNNLQTIRSTYFAHLNKLMDLD      714
AnStephensi_XP035917013       SYNGTFNVYLMDISNCDISELADNTFKTMPQLTRLNLAWNNLQTIRPPVMAHLSKLMDLD      704
IndInt_g22212                 SYNGTFNVYLMDISNCDISELADNTFKTMPQLTRLNLAWNNLQTIRPPVMAHLSKLMDLD      820
                              *:****.**:**:*****************:****************   :***.*****
```

```
AnColuzzii_XP040230463        LSNNMIINMDEKTFQNNRNLNTLNLAGNQIATLTSKLFHPLQYLSELDVSDCDLRTIWED    766
AnArabiensis_XP040158968      LSNNMIINMDEKTFQNNRNLNTLNLAGNQIATLTSKLFHPLQYLSELDVSDCDLRTIWED    766
AnGambiae_AGAP006645          LSNNMIINMDEKTFQNNRNLNTLNLAGNQIATLTSKLFHPLQYLSELDVSDCDLRTIWED    774
AnStephensi_XP035917013       LSNNMITELDEKTFQNNRNLNSLNLSGNQISSLASKIFHPLQYLSELDISDCDLRTIWDD    764
IndInt_g22212                 LSNNMITELDEKTFQNNRNLNSLNLSGNQISSLASKIFHPLQYLSELDISDCDLRTIWDD    880
                              ****** ::************:***:****::*:**:***********:*********:*

AnColuzzii_XP040230463        SAAGTKREEVLPNLKRLNVSYNEIMEVFVSDLESMGKLRVLDIRNNTLTCNERLPTLIDW    826
AnArabiensis_XP040158968      SAAGTKREEVLPNLKRLNVSYNEIMEVFVSDLESMGKLRVLDIRNNTLTCNERLPTLIDW    826
AnGambiae_AGAP006645          SAAGTKREEVLPNLKRLNVSYNEIMEVFVSDLESMGKLRVLDIRNNTLTCNERLPTLIDW    834
AnStephensi_XP035917013       SAVKAKREEVLPNLKRLNASYNEISEVFVSDLASMAKLRVLDIRNNSLSCNGRFPTLIDW    824
IndInt_g22212                 SAVKTKREEVLPNLKRLNASYNEISEVFVSDLASMAKLRVLDIRNNSLSCNGRFPTLIDW    940
                              **. :*************.***** ******* **.**********:*:** *:******

AnColuzzii_XP040230463        LQKKQISMGLGDTHDHVTHAEMTTELRTDNSATLKQWGQFAWEICQGGGSDSPNQRMTYS    886
AnArabiensis_XP040158968      LQKKQISMGLGDTHDHVTHAEMTTELRTDNSATLKQWGQFAWEICQGGGSDSPNQRMTYS    886
AnGambiae_AGAP006645          LQKKQISMGLGDTHDHVTHAEMTTELRTDNSATLKQWGQFAWEICQGGGSDSPNQRLTYS    894
AnStephensi_XP035917013       LQQKQISMAD-NSHDHVTHAEMN-VLRTDNSVSLKQWGQFAWEICQGGSG-DPDNTLSYS    881
IndInt_g22212                 LQQKQISMGD-NSHDHVTHAEMN-VLRTDNSVSLKQWGQFAWEICQGGSG-DPDNTLSYS    997
                              **:*****.  ::*********.  ******.:***************.. .*:: ::**

AnColuzzii_XP040230463        ESDEEYNIEDEDDVSEAADADVMAEAKNAKSNKIVDSKNELDDYSDDSVDSDDSEDEDSA    946
AnArabiensis_XP040158968      ESDEEYNIEDEDDVSEAADADVMAEAKNAKSNKIVDSKNELDDYSDDSVDSDDSEDEDSA    946
AnGambiae_AGAP006645          ESDEEYNIEDEDDVSDAADADVMAEAKNAKSNKIVDSKNELDEYSDDSVDSDDSEDEDSA    954
AnStephensi_XP035917013       ETEEDYDEDEEDDSED-ADAEV-TADTNAKSNKIVDSKNELDEYSDETADSDSDDNDNAD    939
IndInt_g22212                 ETEEDYDEDEEDDSED-ADAEV-TAGTNAKSNKIVDSKNELDEYSDETADS--DDNDNAD    1053
                              *::*:*: ::*** .: ***:* :  .***************:***::.**  .:::::

AnColuzzii_XP040230463        --DEDD---EEDDEEDDDDDDDDEEEQERENILDAANGIQKIEQGILTGKQADED--DDG    999
AnArabiensis_XP040158968      --DEDD---EEDDEEDDDDDDDDEEEQERENILDAANGIQKIEQGILTGKQADED--DDG    999
AnGambiae_AGAP006645          --DEDD---EEDDEEDDDDDDDDEEEQERENILDAANGIQKIEQGILTGKQADED--DDG    1007
AnStephensi_XP035917013       ADSEDDEDDEDADDDDDEDEEEEDDVQERENILDAANGIQQIEQDILTGKLNNNADDDDD    999
IndInt_g22212                 ADSEDDEDDEDADDDDDEDEEEEDDVQERENILDAANGIQQIEQDILTGKLNNNADDDDD    1113
                                .***   *: *::**:*:::::: **************:***.*****  ::   **.

AnColuzzii_XP040230463        EYE--DDGASVDDVVILENGSMFAVSSVWIVVACILFGISMLLLVVVKVLTEVMKRRGER    1057
AnArabiensis_XP040158968      EYE--DDGASVDDVVILENGSMFAVSSVWIVVACILFGISMLLLVVVKVLTEVMKRRGER    1057
AnGambiae_AGAP006645          EYE--DDGASVDDVVILENGSMFAVSSVWIVVACILFGISMLLLVVVKVLTEVMKRRGER    1065
AnStephensi_XP035917013       EYEDDSEGTNVDDVVILENASMFALSTVALMVSCIVIGSLIMVLLVAKIVGLIFKRRGER    1059
IndInt_g22212                 EYEDDSEGTNVDDVVILENASMFALSTVALMVSCIVIGSLIMVLLVAKIVGLIFKRRGER    1173
                              ***  .:*:.*********.****:*:* ::*:**::*  :::*:*.*::  ::******

AnColuzzii_XP040230463        YRQALLASKNSFVYQKLTEDIVPSKVPKIHRYEPINQV    1095
AnArabiensis_XP040158968      YRQALLASKNSFVYQKLTEDIVPSKVPKIHRYEPINQV    1095
AnGambiae_AGAP006645          YRQALLASKNSFVYQKLTEDIVPSKVPKIHRYEPINQV    1103
AnStephensi_XP035917013       YRQALLASKNSFVYQKLTEDIVAPKVPKVHRYEPINQV    1097
IndInt_g22212                 YRQALLASKNSFVYQKLTEDIVAPKVPKVHRYEPINQV    1211
                              **********************  ****:*********
```

*Supplementary figure 4b : MSA for IndInt gene g22212.t1 across other Anopheles species shows high conserved homology.*

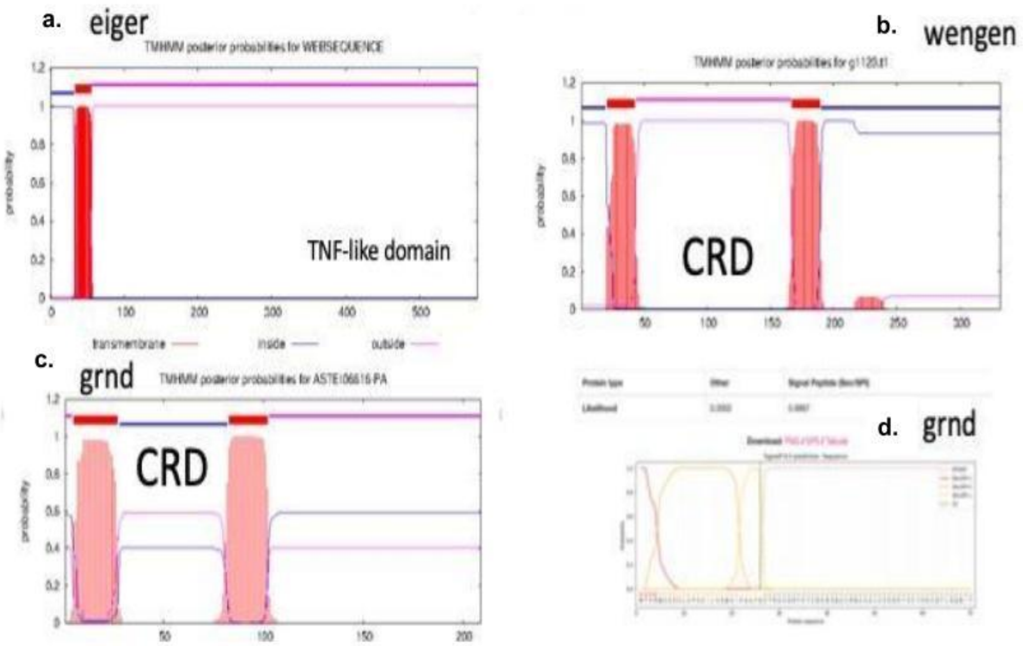


*Supplementary figure 5: (a) transmembrane prediction for eiger (b) transmembrane domain prediction for wengen (c) transmembrane prediction of grnd (d) signal peptide prediction for grnd.*

**UCI**

```
# WEBSEQUENCE Length: 475
# WEBSEQUENCE Number of predicted TMHs:  0
# WEBSEQUENCE Exp number of AAs in TMHs: 0.00271
# WEBSEQUENCE Exp number, first 60 AAs:  0
# WEBSEQUENCE Total prob of N-in:        0.02124
WEBSEQUENCE     TMHMM2.0        outside      1   475
```

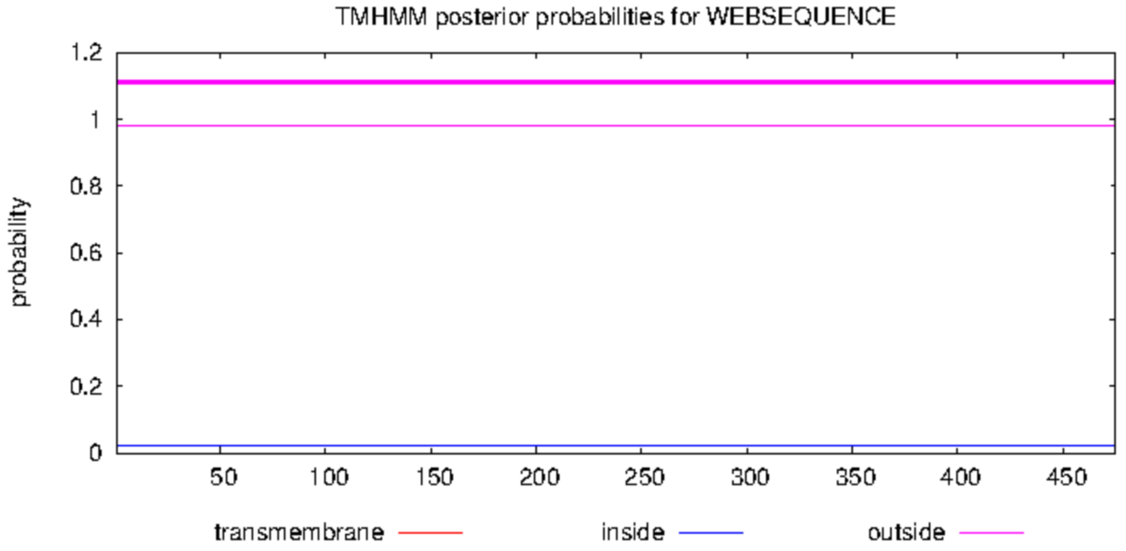


**STE2**

```
# WEBSEQUENCE Length: 455
# WEBSEQUENCE Number of predicted TMHs:  0
# WEBSEQUENCE Exp number of AAs in TMHs: 0.00349
# WEBSEQUENCE Exp number, first 60 AAs:  0
# WEBSEQUENCE Total prob of N-in:        0.02710
WEBSEQUENCE     TMHMM2.0        outside      1   455
```

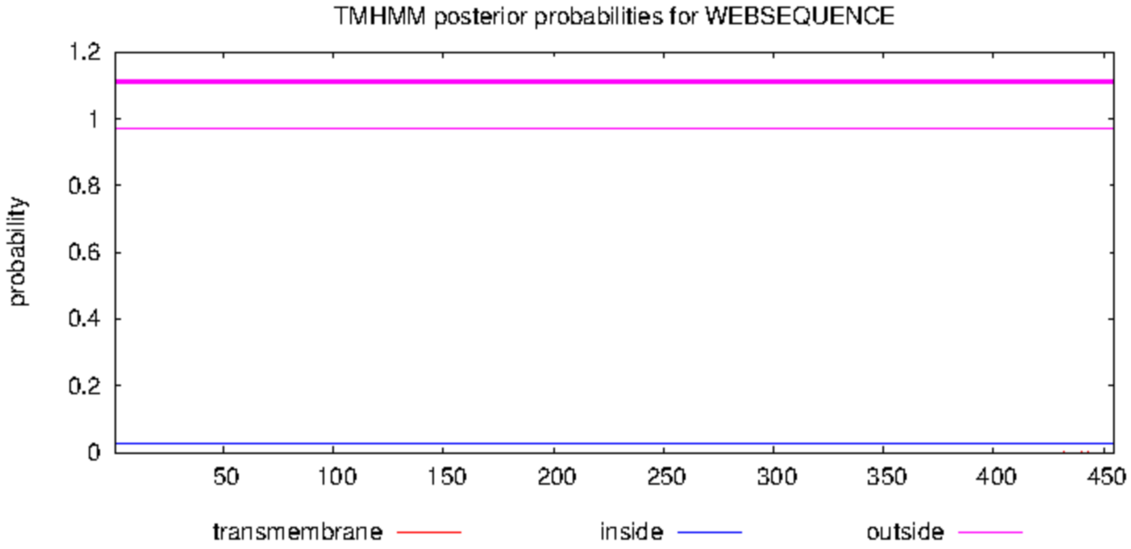

***An. gambiae:***

```
# AGAP006771-PA Length: 544
# AGAP006771-PA Number of predicted TMHs:  1
# AGAP006771-PA Exp number of AAs in TMHs: 22.71772
# AGAP006771-PA Exp number, first 60 AAs:  22.71771
# AGAP006771-PA Total prob of N-in:        0.99203
# AGAP006771-PA POSSIBLE N-term signal sequence
AGAP006771-PA   TMHMM2.0        inside        1    32
AGAP006771-PA   TMHMM2.0        TMhelix      33    55
AGAP006771-PA   TMHMM2.0        outside      56   544
```

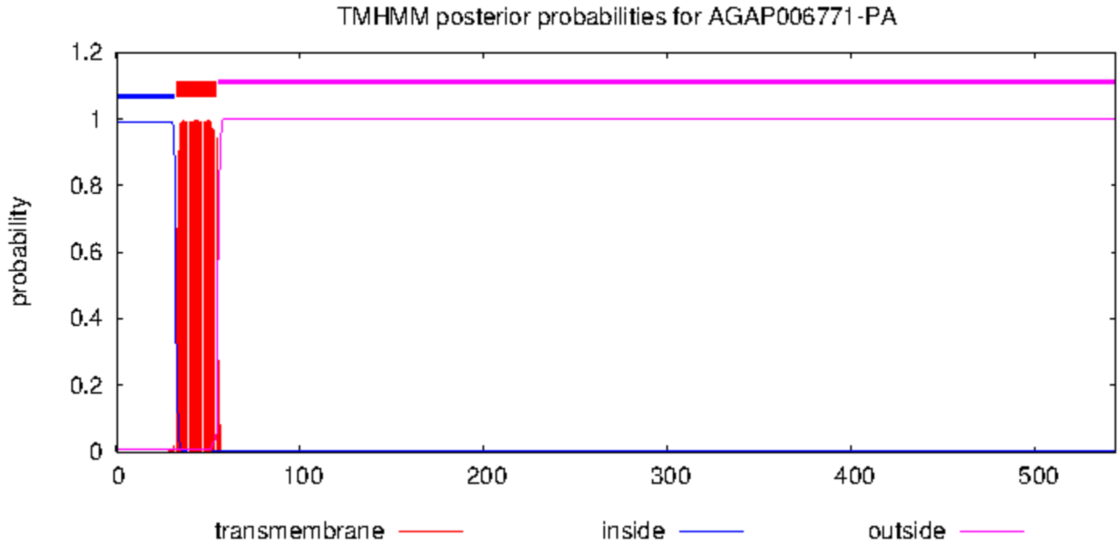


*Supplementary figure 6: TmHMM analysis of the eiger gene in the UCI and STE2 strains and the An. gambiae PEST strain, respectively.*

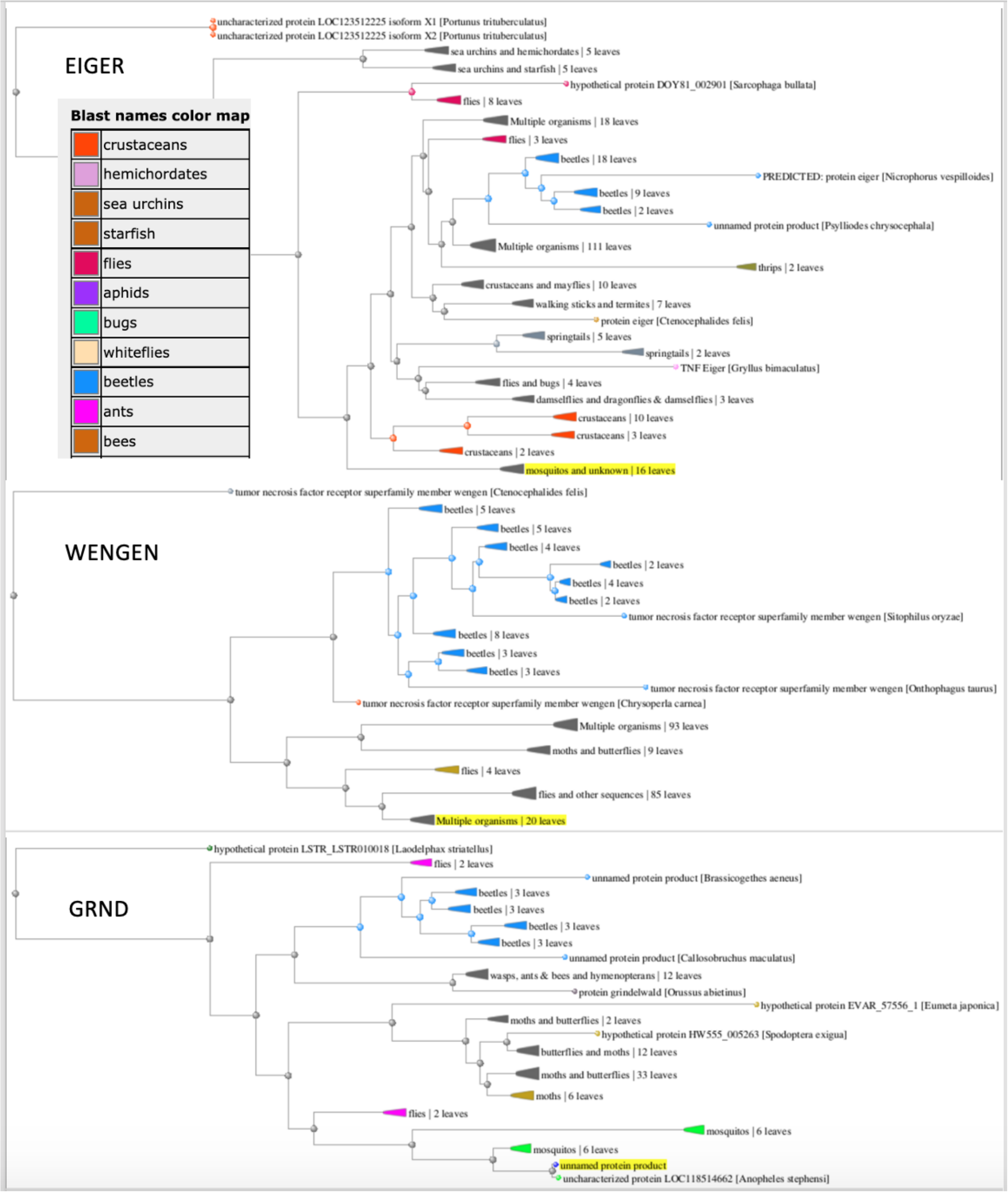


*Supplementary figure 7: BLAST result of eiger-wengen-grnd proteins from IndInt against NR-protein database at NCBI suggesting diversity in eiger gene across vectors across arthopods.*

**************************************************************************

**>ANSTEP-UCI_TRAN_00009302-RA protein Name:"Similar to litaf Lipopolysaccharide-induced tumor necrosis factor-alpha factor homolog (Xenopus tropicalis OX=8364)" AED:0.00 eAED:0.00 QI:437|1|1|1|0|0.33|3|1007|126**
MTKDGPPPYGFVAPPSAPPSYAQAVGGVPPSSPFTPQQPVLTSAQIVTTVVPIGPQSTHMVCPSCH
SEVVTKTTTSPGMIAYVSGFLIALFGCWLGCCLIPCCIDECMDVHHTCPHCKAYLGRHRR

**>ANSTEP-UCI_TRAN_00010133-RB protein Name:"Similar to Litaf Lipopolysaccharide-induced tumor necrosis factor-alpha factor homolog (Mus musculus OX=10090)" AED:0.14 eAED:0.14 QI:206|1|1|1|0.66|0.5|4|598|158**
MNPSGKGSGPEGFQAQPLNQPPYPAQSPPYPGQMPAATGGYPHPSANVPNYAHPPPPPPYDANSNV
IPPPQNAGTTYVQVVTSPQVGPDPASMVCPSCTKHVITRLDYETSTKTHIAAGLLCLFICWPCFWI
PYIIDSCKNANHYCPNCGAYIGTYRG

**>ANSTEP-UCI_TRAN_00010132-RB protein Name:"Similar to Litaf Lipopolysaccharide-induced tumor necrosis factor-alpha factor homolog (Mus musculus OX=10090)" AED:0.01 eAED:0.01 QI:206|0.5|0.66|1|1|1|3|787|124**
MDPPPYDQINRPVPAAYPHQQTPTADPFKSASLHTQEPPSLQTTVIVTSPQVGPDPTTIICPSCRA
TVVTRLEYETTTKTHLCAGLLCLFLCWPCAFVPYCSTACRDANHYCPNCGSFIGTYRK

**>ANSTEP-UCI_TRAN_00010131-RA protein Name:"Similar to FV3-075L Uncharacterized protein 075L (Frog virus 3 (isolate Goorha) OX=654924)" AED:0.00 eAED:0.00 QI:483|1|1|1|1|1|2|936|82**
MTTIIVTNPQVGPDPMTITCPSCRATVRTKVKHESTTSTHACALLLWIVCWPCLCLPYCC
NSCRDANHYCPRCNTFLGSYKH

*Supplementary text 1: FASTA sequences of the LITAF genes from the UCI strain*

**>g6300_ADAM17_TACE_IndInt**
MSLPSIMLTVVLVVSFAVLIVPLQGQLHKNLKYYETLHAKDLSHRIEKRGTKHSTHPFNTIKEVEF
KVLGRKFRLILHPHASVLHSNFRAYSVDGNGSESIVHLDRSNFFKGRVFGEMESHVNAHIDDGVMT
ASVVLPDETYHIEPSWRHLPHLSDKHMIAYRTSDIKFSWDQVDAISGDLGVPRTCGYVKEGLELEG
QPDDGDEAADGGQAFADGGVDAYENDRDPTPETVWHAEDSQDARKSRRKRQADQYEYTPTKTRCPL
LLVADYRFFQEMGGSNTKTTINYLVRAPPPTGSFGAPFVVPFSNRFLPFYQISLIDRVHKIYNDTI
WQDRSDQEGFKGMGFVIKKIVVHSEPTRVRGGEAHYNMVREKWDVRNLLEVFSREYSHKDFCLAHL
FTDLKFEGGILGLAYVGSPRRNSVGGICTPEYFKNGYTLYLNSGLSSSRNHYGQRVITREADLVTA
HEFGHNWGSEHDPDIPECSPSASQGGSFLMYTYSVSGYDVNNKKFSPCSLRSIRKVLQAKSGRCFS
EPEESFCGNLRVEGDEQCDAGLLGTEDNDACCDKNCKLRRNQGAVCSDKNSPCCQNCQYMMAGVKC
REAQYATCEQEARCTGNHAECPKSPPMSDGTMCQERGQCRNGKCVPYCETQGLQSCMCDIIADACK
RCCRQSINETCFPVEPPDVLPDGTPCIQGFCNKGMCEKTIQDVVERFWDIIEEININKVLRFLRDN
IVMAVVMLTALFWIPVSCVIAYFDRKKRKEDWKEYEWSQKLDLIHPSDRRRVIHIRVPRQKITVAR
M

**>g22826_Eiger_IndInt**
MTAETLKPFLNLPNATANDLKAHCAQRSTVKRGLVALGCILAAALCCGLIGVQIWHLNKTAILQQE
VDDLKLQLYLRAREFTDYEASASARPSSLTITTPDRMIMCADGVRRTPVESQRDLFMMGLVDACKN
IQTLVSLICPSFDNELFDSPDEYVYEPEDLDPDVSDDGHREQETNESGLSSSGMQGDDLELDDSEE
KENLGIEDLRETNDDDDDDDDIIIDDRVDIGVPAGGKRRARSISGVTRQGVPIVDEPYVPRRNRTR
HPHRIFEQLRQRPEELVTPPTSAEMFRWDVDSRKTSHTAARHQSYGSISIRPYQHEAHSLREHHST
TPMPVAYRSHGGVRHHSRNELHAGNPTKPPAQILNRMSRVQATGDSMRKQLQQTQRFPKVVGNPGT
QKEIVMAPESRVRLRQRKGGAVQPAEPVAKGVHLVELSTMDAVHSDSRYQEWSANDNASKQAIQSS
SFAFDGERLTVNEPGLYYVYAQVTYSNEFEANGYKVLINGRKHLSCTVNTAQQGENTNTCFTAGLA
EIAHAGTTIAVEDVAHGRQHVMYPEKTFFGAFKIGRLAAPTVTHRRARKVA

**>g1129_Wengen_IndInt**
MTHVPDIRCGMAQVGRRRSHPRTGRGVIPTTTLVLLAILLQLMDLGDGPGGGRSMLVRAACESRKS
WWDPTLGECAPCLICADHQVVLRPCQDYMNTVCGTMKDLTASVSRPYPHLPEGNGNGIGSVGRTRA
HHWKEERRKEGDGVEGYRRVVPAISTEEILWDWQVASLLIAIIGCLLFLLAAGCVALNQSRQWRRI
EKHFDAAQNISKPINNERAPYGILAVGFAVVDMEALSAQLVNHLASMQHLESGPILLENFDHGRLR
TTGHHPIEVRCVYLDQLLDEKCSQKAHSVQPTAGNLYIEESIDTPRLQSPIPSARSPPPPPISLIR
HY

**>g18030_Grnd_IndInt**
MTVRWTSSVLVPVVILVAWSIPSAFGCTRKCDRFEYCDENSNACRNCSALCEKDEYSCYHKCQTLR
QNITTLQESVLTMKTIVLVFSVVAFVLICTALALLIKKYVHCIREFCRWRPQNKNTTTPPVAYTHE
NPNTKSPKTVPKNGANGPKQTSTSVSIYPETEVDHSVQTGTTSISHRYPAEDSTESYSYDNAACNV
TPTSNNPMPKF

**>g5130_TRAF2_IndInt**
MSRPVKREVSAEENCLQSAAKVDDDLLEARYECPICSCWLNEPILTKCGHRFCRKCITDWLNGKNS
ICPLDNEPLDIKCDIFPDNCTRREISQIKKPCPNSIRGCVDQFSPTEIDSHLRQCPFAMSRQSQPC
PFARIKCKFIASDEDALNAHIASDCQQHLQLLLETYTGSNDRYKFWDPPAKNSVPEMSTNELVRSM
YERIVILEQEVHILGIKLSKQELQLTKINQEVDPRYSGGVLLWKLEDFSNKIDSMVANSNCMFYSG
QAYTSPHGYKFCARINVSPRTKDSIGLHVHLMQSENDYHLEWPFKGRIKITLLNVRSPELSQHDTI
MSKPEILAFHRPHQDISPRGFGFLEFAKIKEILAKFADNNTVVLKIQMNIV

**>g13550_TAB2_IndInt**
MCSDLGRYDYQFSIRLRVATAKGAAVRTNCGWLREENERTENRESERSDEAEFDDRTVLRCDGEKK
DDFVSGYQGVWWWFLQTPAIPSTTQSNNTQHYYDEPATAVFTSNSSAATSGSGSNAADSSSILKQR
ATCACSNISIMQLFHEMKQKYPTVPDTVVSELVTQNCHDRPACIGKLEEAVLGTPAQTTYPSQSIH
SGSLKRRSGDRKLAGNRSDGSNSYSSSNSSSNSSRESSVDSSRLQSASYPTTHQQQQQQGVPTTGG
GTGYSDNRFIGESRISSSCANRPTTLAVRPAAAGTGVGNFGATPPPPPMRPNRIAPPCPPNNVPLG
TSTITTTTTASSSTELGETVNLQVNVTVSPKAGGPMIAGQRHTSTISLQPEPPYSRELAQVSSTFS
GGNPLVMGSPTAGTVAGSTPGGSSRTGSASVGIGAGGNGGRSSTSVNLTLRQPTDGRPRTPIHIHA
SPLKYTAKNFNAQSGIQSKLEVTFRDGFGSFSAMRAQVPGYEPQREMMASPQQPEREPCAVALPDG
HSGWYQSPSSAGSRVPYGGLSGFPPPLSSSSPLPNLAAHQMVSSRGMNGQSLFADGQLENDRLTKE
MRYQNFVVAEMAVSQQLEQKQRLSLEVERKRTQFESICREIFVLQQPLRFIDAELLDREVLVLAAE
VEQLQKEVDSCDEEEARIQAAAAAAATGSSITDGLSALSVEGGSPLMLPPSMVSGASGGGTVVNRP
PRPPRPPPPRAPTQSPSRGSSGSVTPSTPHLAPSLAVIGGTSSPVPTPTSSVSSSCSTSSSFQSNG
GGVGDPTAPNRLQPSATTGPNQPWTCSLCTFQNHELMPACEVCSLPKASGSRTAVAAPMTNSNPAF

AVAGDVRDGVGGSPHAAGVLLRRQHHLSVDTGVAAAAAAAAAAAAAAVVGASVAGPSTAPPYPGVA
EMPPPTFASLTTGHTSTSMPLQPIPLSHGQQQQQQQQQQQQPTQTVQQQSQQSNPPPPLQNAQYQK
SSIEC

**>Intsfg5795_Hep_IndInt**
MSDDQSSLSSPATTPSSARPFMPLPLELNGERRKPSKKLSFQGCQGGQAPMIPDPTRERIRMQAAA
GKLQIAPNQTYDFTSDDLLDEGEIGRGAFGAVNRMKFTHTGTVMAVKRIRSTVDEKEQKQLLMDLE
VVMKSNDCNTIVTFYGALFKEGDCWICMELMDTSLDKFYKFICECQQSRIPEPILAQITFATVRAL
NYLKEELNIIHRDVKPSNILLKRNGDIKLCDFGISGQLVDSIARTKDAGCRPYMAPERIDPQRAKG
YDVRSDVWSLGITLMEVATGKFPYPKWGSVFEQLSQVVEGDPPRLCTTYNGMEFSIDFVNFVNTCL
IKEERDRPKYGKLLQHAFIQHAEKSDTDVAAYVSEVLESMANNGITQFTTNLPAEGWNESFN

**>g5319_MKK4_IndInt**
MSDDQSSLSSPATTPSSARPFMPLPLELNGERRKPSKKLSFQGCQGGQAPMIPDPTRERIRMQAAA
GKLQIAPNQTYDFTSDDLLDEGEIGRGAFGAVNRMKFTHTGTVMAVKRIRSTVDEKEQKQLLMDLE
VVMKSNDCNTIVTFYGALFKEGDCWICMELMDTSLDKFYKFICECQQSRIPEPILAQITFATVRAL
NYLKEELNIIHRDVKPSNILLKRNGDIKLCDFGISGQLVDSIARTKDAGCRPYMAPERIDPQRAKG
YDVRSDVWSLGITLMEVATGKFPYPKWGSVFEQLSQVVEGDPPRLCTTYNGMEFSIDFVNFVNTCL
IKEERDRPKYGKLLQHAFIQHAEKSDTDVAAYVSEVLESMANNGITQFTTNLPAEGWNESFN

**>Intsfg11051_PI3K_IndInt**
MKQEQTNTTMKRTAHSTLPGNDPSWQVPHRIYLQQQHQQHLHQQQQQQPMQQRPLFDYEFWKNPDE
LTELHFLMPNGVMITMCIPIHITLEELKTDVWEEAERYPLYCHLGDKNKYCFSTLASGRTNCNTTH
ENEATRLMDVQPALGILRFVERTNVSEDTKLMENISILLDSKLPMKTLVNPEVNDFRAKMSLLSEE
IGSRRAAMTRMERIAYQYPAKLVSGSHVPDQISRRLANVDGHFCVVAISTDMQVTVKVPCRATPDE
VLNRILEKKRISMKARMDNSSDFILKVCGREEYIYGAYPMISFQYVQDCLSRDETPTFVPRLVRSV
EVFKNDIYDARDDFTQWSSTTATTNTTSSALVSTSSSVSTMSLVSVGKHNSSIASTGSSSVYSSSS
QSQQSHTLRKVKYVTSWEVDTKLQCTVQEIRGLNIESDKELGVQLGLFHGGKSLCKTARTRTVTVN
SGKAVWNETICFDINVSNVPRMARLCLVVYENMRTTKSTGIRTRRTKDGLINPIAWVNTMVFDYKN
QLKSDSVTLYTWDYAEDAQSEDILHPLGTVEPNPNRDNNMFVSMLLSFGPYGPEGRIIVYPGEEEL
LAHAAKMSHRNEHLNRESAEDTRSIKAIMSAYMYNDRLNDIHEQDRNAIWAKRRECMLQMPQGLPC
LLYCVEWNNRDEVSEIVSLLQEWPKLLIERALELLDYAYADKYVRRYAVDCLRTIEDDELLLYLLQ
LVQALKHESYLNCDLVYFLLQRALHNQHIGHYLFWHLRSELSVPSVQVRFGLILEAYLLGSPEHVG
VLLKQMQCLRYLQICSDNVKKGSKEKGRALLMEQLAKEGHLTSDLISPLNPSFRCKAVRTERCKVM
DSKMRPLWIVYENSDPNGDDIHMIFKNGDDLRQDMLTLQMLRIMDRIWKSHGFDFRMNPYSCISTD
HKLGLIEVVLNAETIANIQKERGMFSATSPFKKGSLLAWLREHNNTDELLAKAIQEFTLSCAGYCV
ATYVLGVADRHSDNIMVKKTGQLFHIDFGHILGHFKEKFGFRRERVPFVLTHDFVYVINNGRTDRE
AQEFCHFQTLCEEAFLILRQHGCLILSLFSMMISTGLPELSSEKDLNYLRETLVLDKTEEEARTHF
KHKFSEALANSWKTSLNWASHNFSKNNRQ

**>g17575_NIK_Brun_IndInt**
MLSQGPGDPIMAHPDYEQHHYHHGCLLILVRGIGSSKPRSLQRVFERVQRVNNVKIPADSTGTPRD
IWVRYIRDHPVENNDWGDFQTHRRLLGLITVGKFEAQSELNELCRVHESLKVKYTHTLSRCLLFGP
STDELQKLNGAAAGTAVAPDGGSKATLEKCFQTPSNFKSRAFFYPENDPCSNLETKISEFITSIYY
ILELKRMEKTREKLEKAPLLLAPFEKKDFVGLDLESRNNKKRCIGRMTKHLGDLTLQAGLVAESLN
FFHAASETLRAISDSLWLGAANEGLCAASAILLYPNFRYTMSIQRNSSLQENSSSPQKFNLARNQL
YTGSYNGADSSTLKHKKSDMVINLSASDTATILGADKTSASSNSSASSISSSLSSGSGGSGTASGS
SSSDSGTLVNRAGSGVAAKFPSNILQPDEITVRYRDAIINYSKYRNAGIIETEAALKAARICIEQG

KNLDVAMFLQNVLYINLNMTEQQRVRRFEVLTDLYQKIGYNRKAAFCQRLAAWRHVAQSNSNPDWG
QSYRLMLESFSGHKLSLEPNEVLENNMGWPVLQIDLLQQLVGTARRLGQSALATRHMTFLLQTMWK
HLTAQEQREMALQLQNLSAQCEGAPVPLVLENGIVIPPANLTDLPHCSQLLVKDLAPHLKPVKIVV
NKVDSGPFLFTPIHFSSLDRRGIEKDDSKITFNWVQHDVCEVSVSLMNPLPFELQVTDMRLLTTGV
VFEAFPQTVTLQPNVSTNVSLHGTSIECGELEIQGYSTHTLGVKSNCRLKHMLHRRERHLPPCYKV
KVIPALPKLETKTSLPQTATFSGMPNADFVTTSASITLYNGERGECTITLTNSSNIPIEYVDATFH
STLEASLQSRIFQLATDELHRKLPIQPNASIDFKLVIFGEADFLGALTAGPGQTAGYSLPHNPDGM
NSAGPQSLTVSHGGGGGPLGGGMLSAGGGSGNPSIPSRISSPTNTHRRNELLTSSFRSSHSGHSSL
ATLSVGISTGHVPRQLDAQLRFKYSGGEGLQEGFCRQCAISFNVELLPSAQITNWDVLSAEIPSQF
YLVLDVVNLTAQEMSLNYTSNKTILIEAKESCRVPVPVQRCPLERIFAAVAEHQQQHHNHLHQHHH
HHSIGSGSADHHNMPSILSGGAGTSTSDSSDLTERVCSEHISENVNLKWSLPGIDCHGTASLRGIT
LSPAMLDLVTVPPLEWEVKVDDQVVAPQSEVTCVTGQFLSFSISICNLSASVLHQVQLSVQFYQDY
QNGVQNYRLETRVTMSGPNHILIPSLNKDEKAFHKCSVLFFT

**>g8096_AIP1_IndInt**
MHPERNPDALAKLHKNGFEKVEIAVYNVLAREFIYATLPRTQRGQPIVLGGDPKGKNFLYTNGHSV
IIRNIDNPEIADIYTEHSCAVNVAKYSPSGFYIASGDQSGKIRIWDTVNKEHILKNEFQPIGGPIK
DISWSPDSQRIVIVGEGRERFGHVFMAETGTSVGEISGQSKPINSCDFRPARPFRIITGSEDNTIG
VFEGPPFKFKMTKQDHTRFVQAVRYSPSGHLFASAGFDGKVFMYDGTTSELVGEVGSPAHSGGVYG
VAWKPDGTQLLTCSGDKSCKLWDVETRTLISEFPMGSTVDDQQVSCLWQGNHILSVSLSGFINYLD
VNNPTKPLRVVKGHNKPITVLTLSDDRSTIYTGSHDGAVTNWNSGSGTNDRVAGVGHGNQINDIRA
AGDFVYTAGIDDSIKQISVEGNTYTGVDSKLACQPRGMDILKESNTIVVGCVKDITVLQDNRKVSS
VPISYESSSVSINPETMDVAVGGDDSKVHVYTLHEGQLTHKLDLEHLGPVTDVRYSPDNKLLVACD
ANRKVILYSVAEYKPPHNKEWGFHNARVNCVAFSPNSELVASGSLDTTIIIWFVNKPAQHTIIKNA
HPQSQITGLVWLDNETLISTGQDCNTKVWNIENVA

**>g24355_IAP_IndInt**
MDHQYDKGSLLLCLAIFVHKLTIVFHWTQNYLKVLALARVQQRKAGGEVVSSSESAAPSSTPKAPG
DADDDMREMAHALSLPAPIDLPDNKQKDDDLACMSPEYFHIEENRLRTFGRWPVAFISPNVLARYG
FFFVGTDDTVKCYFCRVEIGLWEPQDDVIQEHLRWSPFCPLLKKRPTNNVPLNANYLDAVPEPSYD
TCGISVRQHSYAENVNDRARPDLDRMSGDSWSGASDISLSSSGSSSHNGGDAEPMSGVSGLSSGGG
GVGGGGGLQQDRASMTAAEWNNGVLMGEHSLMRRPEYPNYAIEADRLKTYEDWPTSLKQKPQQLSD
AGFFYTGKSDRVKCFSCGGGLKDWEQDDEPWEQHAIWYSNCHYLQLMKGREFIEKCNELKEAAASA
TSGTSSSMSSASSQPSTSGISSASSSVMSTSPASSSGFSSPTPAADEERTLRCTDHYSSGSDEGGE
DDAGHNRKVPSDGKICKICFVNEYNTAFMPCGHVVACAKCASSVNKCPLCQQPFINVLRLYLS

**>g17640_JNK_IndInt**
MSSRHAFYTVEVGDTKFTILKRYQNLKPIGSGAQGIVCHTKAKDRTKPLTKEPRQGQPPPQQQQQQ
QHTVANMNRRANNYVRVQFGDTEFEVPDRYINLEAKGFGAQGTVCAAYDTVTQQNVAIKKLSRPFQ
NVTHAKRAYREFKLMKLVNHKNIIGLLNAFTPQRTLEEFQDVYLVMELMDANLCQVIQMDLDHERM
SYLLYQMLCGIKHLHSAGIIHRDLKPSNIVVKSDCTLKILDFGLARTAGTTFMMTPYVVTRYYRAP
EVILGMGYKENVDIWSVGCIMGEMIRGGVLFPGTDHIDQWNKIIEQLGTPSQSFMARLQPTVRNYV
ENRPRYTGYPFDRLFPDVLFPTDSNEHNRLKASQARDLLSRMLVVDPEHRISVDQALVHSYINVWY
DESEVNAPAPGPYDHSVDEREHTVEQWKELIYQEVMEYEARNNLADATDEAMAAAAEAAEASSTTA
AAAAAAEDLQPPSEEPPDAMSPAEE

**>g11102_p38_Bsk_IndInt**

MANFYRTEINKTEWEVPDKYQALTPVGSGAYGQVCSAMDTVHNVKVAIKKLARPFQSAVHAKRTYR
ELRMLKHMNHENIIGLLDVFHPGDNKLESFQQVYLVTHLMGADLNNIIRTQRLSDEHVQFLVYQIL
RGLKYIHSAGIIHRDLKPSNIAVNEDCELKILDFGLARPTENEMTGYVATRWYRAPEIMLNWMHYN
QTVDIWSVGCIMAELLTGRTLFPGTDHIHQLNLIMEILGTPNDEFMAKISSESARHYIKSLPKTEK
RNFSDVFRGANPLAIDLLEKMLELDADKRITAEQALAHPYLEKYADPSDEPTSSLYDQSFEDMDLP
VERWKELVFKEVLNFVPQQHAHIGGEPQA

**>g21984_AKT_IndInt**
MTATDNTVRFSDHTLDKATKAKVTLENYYSNLITQHGERKQRQAKLEASLKDETLSESQRQEKRMQ
HAQKETEFLRLKRSRLGVEDFEALKVIGRGAFGEVRLVQKKDTGHVYAMKVLRKADMLEKEQVAHV
RAERDVLVEADHQWVVKMYYSFQDSVNLYLIMEFLPGGDMMTLLMKKDTLSEECTQFYIVETALAI
DSIHRLGFIHRDIKPDNLLLDARGHLKLSDFGLCTGLKKSHRTDFYRDLSQAKPSDFIGTCASPMD
SKRRAESWKRNRRALAYSTVGTPDYIAPEVFLQTGYGPACDWWSLGVIMYEMLMGYPPFCSDNPQD
TYRKVMNWRETLIFPPETPISEEARDTIVKFCCEAERRLGSQRGIEDLKLVQFFRGVDWEHIRERP
AAIPVEVRSIDDTSNFDEFPDVALEIPAHPQPEGEVLKDWVFINYTFRRFESLTQRARAADTEAKQ
QQQQHIVSPNPQSIDRPTNGMMAQDRVTG

**>g17088_IKK_IndInt**
MSGLHSATFIVRGCAVNRLNIAQPAAFFRYIKTQLSVLCALLQESNRGVSLLVNVLANMIKPFEDP
PLIGDWCREKRLGNGGFGVVSLWRNKQTQQAVAIKKFHILQDRSEITDKHCERWRNEVKLMTETVQ
NENIVRTVNVQPTSFIQELLRSSANGLPVLCMEYCEGGDLRRVLNRVENCSGLREQDVRDVLRSLR
NAVAYLHSLKITHRDIKPENLVLKQQGERFIYKADLGYAKALDKQSLNASLVGTVEYIAPDLIYCD
RYNCSVDYWSMGVIGYEIITGVRPFIPHAPITRWMMHVQQKKSADIAITEDNRENYTYHTDIFPEN
HISDCLRKELESWFRLALEWNPKKRGYVPYTKAEPSLANGENGTKGVKQTEDNKPSTVLKIFSLLD
QILEKRILVLFSLYDCRWIDLEVTPETGMETLRDHVYRVTGIPVDDIEFVLPLEQKQPAVGNDTRP
YDLYLPDFYGKPMVYVVHRGSRDSIVQRDLKPRIPKSITDVFQNIKVKLKPHMLRQFIANSYYFIA
QEQRLYGQVLDGIRNYGLMLNDNIARRKDEIGRMNKIVYAILGGVEYHKLTVCHARDALNVERRIP
SATFEMASKRWIENGARIESNVRKLAEMADTITKRYESVLKRSRDALRHALLHPQLDQHDTFGLRN
VECRYEQTRARLMEKILNEKSHMDMSQAVYECLKQRDVLLRELGFLELQQQILDVRREIQEIEKVV
CKVIDVTEKYKRDLARLKLEHQDEVWKMLSDYGRSAAAGSCENGVVHLDNILPLGVAHTDPPVINK
PNFLVGGPTTPQLQSTVSNESFDANQVCCLGYTDPNVEDLIAANETLIHTTTDLLSNSFSLLKLAD
GAE

**>g13192_Caspase1_IndInt**
MEEFALSASLRHSTDELDANAVNRSTNSAIPSALSTSNDVTAQGSDHYDTSHPRRGIALIINQVNF
KDLRKRDGSDKDRDCISAALQGIGFEIRTLDDPNRKELLAMLEKVANEDHSQHDCLVVVVMTHGKE
NNFLYARDKSYEADRLWEPFLGNACPSLRGKPKLFFVQACRGKQLDQGVRLVNMSIDENVSVDSVP
EPVSYVIPTMADLLVMYSTYDGHYSWRNPVNGSWFIQSLSIVLTATAHREELLHILTAVSRRVAFH
YQSDVPQNVKMDAMKQMPCIVSMLTKLLYFPKKS

**>g22865_NFkB_IndInt**
MSLDHCSIALDVLLLQLQQQQQQQQPPPRLDVPYSLQTEQQQLLGASPNQYTILSMDAPSAAAA
VVSVGDYTLGSSRSYASALSPSSSSASPSSPSSVASPNSRASNMSPQSSASDHSASYTLQNLNLSS
STSSMGYPGMSYQQQQQHQPQHQQHHQQQHQQHHQQQHQQQIQPQQQHYYAPQLLNLDQEHLQTAS
FSYTTSSNEAFASPDADYGKPHLVILEQPVDKFRFRYQSEMHGTHGSLMGSRTEKSKKTFPTVELR
GYSGEAKVRCSLYQVDPQRRAPHSHHLVIKSGELDLIDPHDLDVGGPTGESSDGGGGGGGGGSYVA
TFQGMGIIHTAKKFIAEELYKKLRKHRLCELNREPTEREEQQMQKDATVMARTMNLNQVCLCFQAY
RVEPGTGRWVPICEPVYSNAINNMKSALTGELKICRLSTTVSGVDGGEEVFMFVEKVCKNNIKIRF

YELDEYDQEVWQEMAIFSEADVHHQYAIAFKTPPYRNKDITEPVEVHMQLYRPRDRCQSEPVLFKY
KPRSGMMVPGSSGSGSRKRLRISSGNISSEIPTVIQNDPNAGGGGGGGGVPGAGGGGGAGGGGAGS
TTRLPPLHQPFPMLANHDGSIPEGHEPSSAATINAGTATGHQQDIMSGIGSVTTISKELTKASIIQ
EILNIPTTIASDVAFDSSDFPCNSEEFNKLIQEIGNQQDLVKLETDSEAAAGDTTESVLGRAIADL
VASGDDARQGEMLRKLLALIKLFAGDVNRSRQLLASHWTAANQQQLNCLHAAIRRNDTTIACKLIE
LLDEFRLTDELLDLPNDRNETALHLAVSTNNESIVRELLKAGAKLTFCDYRGNTALHRAVVENVPA
MVRLLLRHGQAGRSRLDCTNDDGLTALQAAVYARNLKITRILIDAGASVREKDLKHGNNILHIAVD
NDSLDIVNYVLEHVNRELGLEQNNAGYTPLQLADAKSATGQGNNKLIVRELLRHFPDGLEKRTAGD
DSQEQEEDIDEDEEEEEEEEEDEEEREGEAIEQDKEQKNETVLDSINLNCDRVNIARLLEEHEPEA
EPQRKATKRSDATTATERKEPAAASAPLFDDQCMEELCELLSGSETWRALGSLLDFHPFFTVWEQG
PSPARMLLGYYEMQQLNMDRLIDMLRALELRDCIRSIDEMICRRMK

**>g376_MKK3_IndInt**
MYGSVSPDLLLIVLRFLLGAARDAVPFWRFGLVWTFTLYDDDVQVKDVFSQRSTGELSLEAHGAGP
GLKWHDQTLAATSTNRGEQEEKEGGSQQPTNNNNTNEARKVFLLSVGVSLVWAVAAAEYRLHCRAT
PPPPSPSAPKMPRKNKPIIKLKLDGDTGGGSTVPDSAEGATPRNLDKRGTLTVGERSFVVEADHLR
KLADLGRGAYGIVEKMLHQPSGTVMAVKRITVTSGMGAGGGLGGGGVQSQEQKRLLMDLDVSMRAS
DCRHTVQFYGALFREGDVWICMEVMDTSVDKFYPVVYKRPGRTIPERILGRIALAIVRALHYLHTE
LRVIHRDVKPSNVLMNRRGEVKMCDFGISGYLVDSVAKTIDAGCKPYMAPERIDPSSCSRTAGYDI
KSDVWSLGITMVEIATGRFPYATWRTPFEQLKQVVNDEPPRLPKSAAPDSEEFSAEFHAFTASCLQ
KKFQQRGNYEQLLAMEFLQRYGAGDVQADTSDMAEYVCAILDERDGVPAAEDGGGELAN

**>g20745_JUN_IndInt**
MRNNVLNSKHTNNNTMESFYEDNNAQFVPSTTTTNGVTTGNGNGLKRPATLELNLAPGKARKTRYN
ASVTMPSVLPSPDMQMLKLVSPELEKIISTNATLPTPTPSAIIFPPSATSEQQQFAKGFEDALLSI
HKKDTTSKLNTNNNNNNNNSNTSNNNNGPSANIGGAPAERLCPPTTAAPTTVVPVVGGTTTNNLAST
VLASGHNNGMSGGEITYTNLDYPGVVKEEPQATSSNQSPPVSPIDMESQERIKLERKRLRNRVAAS
KCRRRKLERISKLEERVKELKTQNSELSGVVCSLKQHIFQLKQQVIEHHNNGCTITLVGKF

**>g7769_RAC1_IndInt**
MSSGRPIKCVVVGDGTVGKTCMLISYTTDSFPGEYVPTVFDNYSAPMVVDGVQVSLGLWDTAGQED
YDRLRPLSYPQTDVFLICYSVASPSSFENVTSKWYPEIKHHCPDAPIILVGTKIDLREDRETISVL
AEQGLSALKREQGQKLANKIRAVKYMECSALTQRGLKQVFDEAVRAVLRPEPLKRRQRKCVVIEMQ
TMASGVAGNAEKETATINPREACKVNNEATKRLTNVGKS

**>g4935_MAPK_IndInt**
MSMNEGIAVVTKVVSTSVRRAPEPTAVLTREDFNAGPISLDEIEPGLWLGNASAAADMGTLEKHTI
RSILSIDSVPLPVHITDHPNLRVRHIQAADVPREDLIRHFEDSNRFIVDSLAEGRHVLVHCYFGVS
RSATIVIAYVMQKYRLNYEAAFQRVKAKRIFVMPNPGFVNQLKLYGRMAYRIDRTNERYKLFRLRL
AGDNVRKAKRLPTECMDVVKTDPGVTQESPEPYVYRCRKCRRVVASRSNLLLHKPKSATLAQSPAK
VGREPEATLEEEGTSDEAECDGQPQMNADGENVADHVTHVQTAVATGSSPSSGGGMNELAEQVRRS
SISSEHSNRSSEKDTPMCNKIFFIEPLAWMTDIYRNTQGRLYCPKCTVKLGSFNWVMATKCPCGAE
IYPAFYLVPSKTEYSTVVQNVQVTV

**>Intsfg7581_Erk_IndInt**
MAEAASSGAAAAAAAASAAGPAAGSTSATNPNAEVIRGQVFEVGPRYTNLAYIGEGAYGMVVSATD
TVTKTKVAIKKISPFEHQTYCQRTLREIKILTRFKHENRLSNDHICYFLYQILRGLKYIHSANVLH
RDLKPSNLLLNTTCDLKICDFGLARVADPEHDHTGFLTEYVATRWYRAPEIMLNSKGYTKSIDIWS

```
VGCILAEMLSNRPIFPGKHYLDQLNHILGVLGSPSQEDLECIINEKARSYLQSLPYKPKVPWSRLF
PNADLNALDLLGKMLTFNPHNRISPVAEEPFRIAMELDDLPKEMLKRLIFDETLRFNHNDNNLPGG
M
```

**>g14460_TRAF4_IndInt**

```
MKFPLLVRLCPGVAPSHDHLGSVSPRSKILGALVPAVCELAVRKSDVIYPDPESEKAIMGSLVFCI
HHKQGCKWSDELRKLKAHLNTCKHDAIQCPNKCGSQIPRVMMTDHLAFTCILRRAICEFCNVEFTG
IGLEEHAGTCSSEPIYCESKCGTRVLRGRMSIHRAKDCAKRLRRCPHCGREFSADTLAAHGATCPR
SPIPCPQRCDAGPFARADLDAHLRDECKALTVPCTFKEAGCRFKGPRHLLEAHLETNTSAHLSLMV
ALSGRQGQQINMLKSAMAKLSTNYTGTLLWKITDWSVKMQEAKTKDGLELVSPPFYTSQYGYKLQA
SMFLNGNGPGEGTHVSVYIKVLPGEYDALLKWPFSHSVTFTLFEQGTLGSGGGGQGGVAESFVPDP
TWENFQRPSTEPDALGFGFPRFVSHELLNRRPFVRDDTVFLRVKVDPSKIVAV
```

**>g488_FOS_kayak_IndInt**

```
MRSVRGSTLYGLGIIAFAWLAPATHHNIHKAVQGACPLAVAWWRFASFDGIGVHSGVPTRTTPTLT
PTTLKNIEQTFMEATNAQINLPYQAGFVPPSYPMEDHSQDGGVSQESLSSNSNGSWAVSGSHGYTG
DDHDSKSSASLEQPSSGTRSRRSATGTDENAKNETVTFAIGTGGSGTGARTGGRRNVGGRRPHKPS
NLTPEEEEKRRIRRERNKQAAARCRRRREDHTNELVDETDQLEKKRQSLAQEIQQLQQEKDDLEFL
LETHREHCRLQARRSPIDLKPVDLDTGGFENTGTFVLPKIKTEPEDEFAAAAAAAAAATQQHHQQP
PPNQLQEQLAIEAAEHGTISKKLKLSHAHSDSLETPTPTSSAFAPLAGVGLAGNSGNKLSATPAGS
NGSRPTRPDSLDVKSAAQYPLIARGDAAGSLAISTPSSGLFNFDSLMEGGTGLTPIAAPISFPPNR
NPLELITPTSTEPSKLCSL
```

**>g17088_IKBK_IndInt**

```
MSGLHSATFIVRGCAVNRLNIAQPAAFFRYIKTQLSVLCALLQESNRGVSLLVNVLANMIKPFEDP
PLIGDWCREKRLGNGGFGVVSLWRNKQTQQAVAIKKFHILQDRSEITDKHCERWRNEVKLMTETVQ
NENIVRTVNVQPTSFIQELLRSSANGLPVLCMEYCEGGDLRRVLNRVENCSGLREQDVRDVLRSLR
NAVAYLHSLKITHRDIKPENLVLKQQGERFIYKADLGYAKALDKQSLNASLVGTVEYIAPDLIYCD
RYNCSVDYWSMGVIGYEIITGVRPFIPHAPITRWMMHVQQKKSADIAITEDNRENYTYHTDIFPEN
HISDCLRKELESWFRLALEWNPKKRGYVPYTKAEPSLANGENGTKGVKQTEDNKPSTVLKIFSLLD
QILEKRILVLFSLYDCRWIDLEVTPETGMETLRDHVYRVTGIPVDDIEFVLPLEQKQPAVGNDTRP
YDLYLPDFYGKPMVYVVHRGSRDSIVQRDLKPRIPKSITDVFQNIKVKLKPHMLRQFIANSYYFIA
QEQRLYGQVLDGIRNYGLMLNDNIARRKDEIGRMNKIVYAILGGVEYHKLTVCHARDALNVERRIP
SATFEMASKRWIENGARIESNVRKLAEMADTITKRYESVLKRSRDALRHALLHPQLDQHDTFGLRN
VECRYEQTRARLMEKILNEKSHMDMSQAVYECLKQRDVLLRELGFLELQQQILDVRREIQEIEKVV
CKVIDVTEKYKRDLARLKLEHQDEVWKMLSDYGRSAAAGSCENGVVHLDNILPLGVAHTDPPVINK
PNFLVGGPTTPQLQSTVSNESFDANQVCCLGYTDPNVEDLIAANETLIHTTTDLLSNSFSLLKLAD
GAE
```

*Supplementary Text 2 : FASTA sequences of the genes from the IndInt strain involved in eiger mediated TNF pathway.*
